# Recycling old drugs: cardiac glycosides as therapeutics to target barrier inflammation of the vasculature, meninges and choroid plexus

**DOI:** 10.1101/2020.04.15.043588

**Authors:** Deidre Jansson, Victor Birger Dieriks, Justin Rustenhoven, Leon C.D. Smyth, Emma Scotter, Miranda Aalderink, Sheryl Feng, Rebecca Johnson, Patrick Schweder, Edward Mee, Peter Heppner, Clinton Turner, Maurice Curtis, Richard Faull, Mike Dragunow

## Abstract

Neuroinflammation is a key component of virtually all neurodegenerative diseases; preceding neuronal loss and associating directly with cognitive impairment. Neuroinflammatory signals can originate and be amplified at barrier tissues such as brain vasculature, surrounding meninges and the choroid plexus. We designed a high-throughput screening system to target inflammation in cells of the blood-brain barrier (primary human pericytes and endothelia) and microglia enabling us to target human disease-specific inflammatory modifiers. Screening an FDA-approved drug library we identified digoxin and lanatoside C, members of the cardiac glycoside family as inflammatory modulating drugs that work in blood-brain barrier cells. A novel *ex vivo* assay of leptomeningeal and choroid plexus explants further confirmed that these drugs maintain their function in 3D cultures of brain border tissues. While current therapeutic strategies for the treatment of neurodegenerative diseases are missing the mark in terms of targets, efficacy and translatability, our innovative approach using *in vitro* and *ex vivo* human barrier cells and tissues to target neuroinflammatory pathways is a step forward in drug development and testing, and brings us closer to translatable treatments for human neurodegenerative disease.

**One Sentence Summary:** We have identified cardiac glycosides as powerful regulators of neuroinflammatory pathways in brain-barrier tissues such as vasculature, meninges and choroid plexus.

## Background

Dysregulated chronic inflammatory responses occur early in disease progression, contribute to blood-brain barrier dysfunction and can precipitate neurodegeneration (*1-5*). Accumulating evidence suggests an inflammatory contribution by cells of the neurovasculature, meningeal compartments and choroid plexus (CP) by facilitating the communication of inflammatory signals from the periphery and within the CNS (*6, 7*). Since neuroinflammation in the central nervous system (CNS) can precede and exacerbate neuronal loss, attenuation of these responses in contributing cell types prior to overt deterioration should be more appropriately explored as a therapeutic option (*8-10*).

Neuroinflammation can be most easily targeted anatomically at several sites; the cerebrovasculature that reaches the entirety of the CNS bordered luminally by the blood, and abluminally by the cerebrospinal fluid (CSF), the meninges which surround and envelop the brain and spinal cord directly communicating with lymphatic vessels, and the CP that produces CSF and facilitates fluid movement by way of glymphatic flux (*11-13*). It has been suggested that the brain’s immune functions have been concentrated at these border tissues, which may be more sensitive to inflammatory cues than the rest of the CNS (*14-16*). We have designed our study to test inflammatory modulating drugs on the spectrum of cell types that make up these structures.

At the cellular level the neurovascular unit (NVU) is formed by the specialized arrangement of endothelia, pericytes, astrocyte endfeet, and neurons that work together to maintain brain homeostasis. We have demonstrated that pericytes derived from post-mortem human brains display a robust inflammatory response when stimulated by classical inflammatory mediators such as tumor necrosis factor α, interferon γ (IFNγ), interleukin 1β (IL-1β) and lipopolysaccharide (LPS), and are likely to be targets of neuroinflammation in both an autocrine and paracrine fashion (*17, 18*). Importantly, endothelia and pericytes in the NVU contribute to neuroinflammatory responses by chemokine secretion, adhesion molecule-mediated leucocyte extravasation, and damage-associated molecular patterns (DAMPS) and pathogen associated molecular patterns (PAMPS) receptor expression (*11, 19-22*). Attenuating the infiltration of peripheral immune cells into the brain or preventing inflammatory propagation by barrier cells such as neurovascular cells is likely to be beneficial in limiting CNS immune responses, although few CNS therapies have been based upon this approach. Namely the MS drug natalizumab that targets immune cell entry into the CNS is a good example of this approach (*23*). In this study in addition to primary human brain neurovascular cells such as pericytes, endothelia cells and microglia, we also made use of explants from barrier tissues such as meninges and CP which are directly involved in inflammation.

The human meninges are composed of three membranes (dura, arachnoid, and pia mater) that envelop the brain and spinal cord. The arachnoid and pia mater, collectively termed the leptomeninges, form a semi-permeable membrane to the CSF which fills the subarachnoid space. The leptomeninges are comprised of several cell types, including macrophages, dendritic cells, mast cells, and fibroblasts and are permeated by leptomeningeal arteries (*24, 25*). Furthermore, diverse leucocyte populations reside in the subarachnoid space providing a conduit for immune-brain communication (*26-28*). Meningeal inflammation often precedes inflammation in the CNS and is present in neurodegenerative diseases, chronic inflammatory conditions, and acute pathogen introduction (*29-31*). Importantly, meningeal-derived factors can permeate the brain parenchyma, presumably through glymphatic influx, suggesting that modulating meningeal-derived inflammation also represents an appropriate target to prevent inflammatory-mediated CNS insults (*32, 33*).

The CP is a highly vascularised tissue that harbours fenestrated capillaries surrounded by apical facing, CSF-producing epithelial cells and serves as a gateway for immune cells from the circulatory system to the CNS (*34*). The fact that the CP is a sink for immune cells is of particular importance when it comes to immune cell trafficking in both sterile inflammatory and chronic inflammatory conditions (*35, 36*). Similarly, the CP itself increases secretion of inflammatory messenger molecules into the CSF with age and disease, which are transported throughout the brain through CSF circulation (*37, 38*). Together this supports the examination of the CP as a target for reducing neuroinflammation.

Here, we utilise primary dissociated human brain pericytes and endothelia, as well as explants from the meninges and the CP to screen drugs for anti-inflammatory properties. Drug screening approaches are not typically performed using primary human brain cells due to limited tissue yields, difficulties of human brain cell culture and accessibility. However, rodent immune cells display numerous discrepancies compared to their human counterparts, with respect to both immune functions and pharmacological responses (*32, 39*), necessitating the use of primary human brain cells for pre-clinical compound screening with an intent for translational drug discovery. Primary human brain pericytes are suitable for high throughput drug screening for candidate compounds with anti-inflammatory functions because unlike many other human brain cell types, pericytes undergo rapid proliferation *in vitro*, allowing for efficient bulking of these cells for the purposes of high-throughput drug screening. In this study, selected hits from initial screens were validated in both pericyte and endothelial cultures to explore the anti-inflammatory efficacy of these compounds in NVU-associated cells. Next we describe and characterise the inflammatory contribution of an *ex vivo* model of human leptomeningeal and CP explants using it to investigate lead anti-inflammatory compounds in a complex multicellular system more closely recapitulating the *in vivo* human brain environment. Finally, we demonstrate the efficacy of two anti-inflammatory compounds digoxin and lanatoside C, in attenuating meningeal inflammatory responses and preventing meningeal-mediated inflammatory propagation.

## Results

### Identifying cardiac glycosides in a screen using primary pericytes

Primary human brain pericytes respond to inflammatory stimuli to induce the expression of chemokines and adhesion molecules, including chemokine (C-C motif) ligand 2 (CCL2) and intercellular adhesion molecule-1 (ICAM-1), respectively (*17, 40-42*). Due to their involvement in enhancing leucocyte infiltration into the brain, these mediators were selected as candidate proteins to determine the anti-inflammatory potential of the Prestwick compound library (*17*). Concentration response curves of IL-1β-induced CCL2 and ICAM-1 protein expression, as determined by immunocytochemistry, revealed an EC_50_ of 0.02 ng/mL for CCL2 and 0.03 ng/mL for ICAM-1, thus 0.05 ng/mL was selected as an optimal concentration of IL-1β to determine immune modulation, allowing for identification of compounds which may induce and attenuate inflammatory responses (Fig. 1A and B). Pericytes were screened for modulators of IL-1β-induced CCL2 and ICAM-1 expression by pre-treating cells with > 1200 FDA approved compounds at 10 µM in duplicate for 24 hours, then stimulating with IL-1β (0.05 ng/mL) for 24 hours. CCL2 and ICAM-1 intensity values normalized to total cell counts confirmed inhibitory effects of transforming growth factor receptor β (TGFβ_1_), a known inhibitor of chemokine and adhesion molecule expression in pericytes (*42*) (Fig. 1C and D). Whilst some plate variability in terms of cell numbers and absolute intensities was observed, the IL-1β induction of CCL2 or ICAM-1, and the TGFβ_1_-dependent inhibition was consistent across all test plates (fig. S1). A total of 82 compounds that met cut-off criteria (changes in intensity ± 20 % over vehicle + IL-1β values, < 50 % cell loss, with standard deviations of < 15 % for either CCL2, ICAM-1, or both) were selected for further validation. Drug classification revealed hit compounds that altered CCL2 or ICAM-1 were mainly comprised of therapeutics categorised as anti-bacterial, anti-asthmatic, anti-helminthic, cardiotonic, bronchodilator, anti-septic, anti-neoplastic, anti-amoebic, and anti-inflammatory (Fig. 1E and F). A second confirmatory screen of the 82 hits generated 44 that had reproducible effects. These 44 hits were then screened at three concentrations (0.1, 1, and 10 µM) for the ability to modify IL-1β-induced CCL2 or ICAM-1 in pericytes, without causing significant cell loss (Trial 3, fig S1). IL-1β-independent effects were assessed by screening compounds in pericytes with the same paradigm (24 hours of compounds at 10 µM, 24 hours of vehicle only for IL-1β), which revealed no significant changes relating to CCL2 or ICAM-1 expression (fig. S1).

**Fig. 1:**
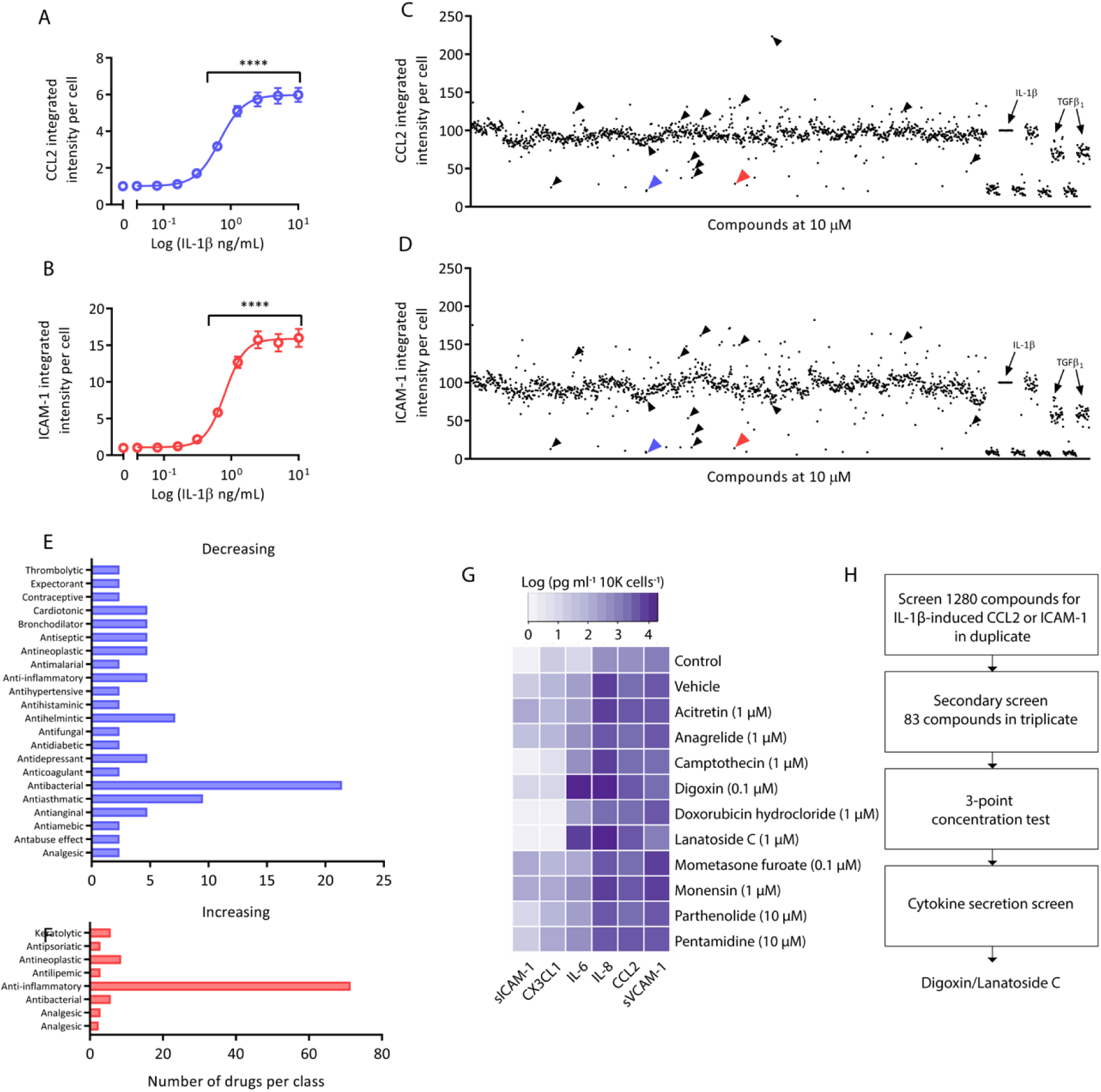
High throughput screening of pericytes identifies inflammatory-modulating compounds. (A-B) Pericytes were treated with IL-1β for 24 hours and immunostained for CCL2 and ICAM-1 protein expression. Integrated intensity per cell was calculated using total cells (Hoechst), n = 3, significance determined using one-way ANOVA with Dunnett’s correction for multiple comparisons, \*\*\*\**p*<0.0001. (C-D) Pericytes were screened for compounds that can modify IL-1β-induced CCL2 or ICAM-1 expression using the Prestwick chemical library. Integrated intensity per cell was calculated using total cell counts and normalized to IL-1β + vehicle condition (arrow)(blue arrow-digoxin, red arrow-lanatoside C, black arrows-hit compounds tested in (G)). Internal controls were present on each drug plate (TGFβ_1_, 1 or 10 ng/mL). E-F). Hits identified using cut-off criteria from primary screen that modified CCL2 or ICAM-1 expression with known therapeutic uses. (G) Conditioned media from pericytes pre-treated with hit compounds for 24 hours, then stimulated with IL-1β (0.05 ng/mL) for 24 hours was analysed by CBA, data was normalized to cell number and presented as the logged value of cytokine secretion in pg/mL/10,000 cells. Control (vehicle for IL-1β (0.01% BSA in PBS), IL-1β (0.05 ng/mL + dimethylsulfoxide (DMSO))(n = 2, media pooled from triplicate wells for analysis). (H) Drug screening pipeline in primary human brain pericytes. Initial screen in pericytes completed in duplicate wells and secondary screen in 83 compounds that met cut-off criteria was completed in triplicate wells. Effects that were reproduced were examined using a 3 point concentration test, then tested for effects on cell secretion (data in fig. S1).

Hits were narrowed down to 10 compounds that consistently modified either IL-1β-induced CCL2 or ICAM-1 expression by immunocytochemistry without significant cell reduction (as assessed by total cell counts). To further investigate the anti-inflammatory ability of hit compounds, their effect on secretions of a larger panel of inflammatory chemokines, cytokines, and adhesion molecules previously identified to be secreted by pericytes in response to IL-1β was determined by cytometric bead array (CBA) (*43*). Concentrations determined from Trial 3 were used to treat pericytes as above (fig. S1). Analysis of cytokine secretion showed that both cardiac glycosides digoxin and lanatoside C had the most substantial effects on soluble ICAM-1 (sICAM-1) and fractalkine (CX3CL1) (a known microglial ligand) (*44*). In particular, digoxin inhibited IL-1β-induced secretion of CCL2, sICAM-1, soluble vascular cell adhesion molecule-1 (sVCAM-1), and CX3CL1 but increased secretion of interleukin-6 (IL-6) more than 6-fold over IL-1β alone (Fig. 1G, fig S2). A generalized pipeline for refinement of the hit compound list leading to digoxin as a lead compound is presented in Fig. 1H. Digoxin is indicated for the treatment of symptomatic heart failure, and atrial fibrillation, thus we were surprised to see anti-inflammatory effects on vascular cells. Lanatoside C, another cardiac glycoside found in the Prestwick library, demonstrates structural similarity with digoxin (*45*). Little is known concerning inflammatory outputs by cardiac glycosides on brain vascular cells, and due to the ability of digoxin to most effectively limit pericyte-mediated inflammatory responses, digoxin and the closely-related lanatoside C where chosen as lead compounds for further analysis.

### Anti-inflammatory effects of digoxin and lanatoside C in primary human brain pericytes

Pericytes express several proteins that facilitate leukocyte extravasation and polarise surrounding immunologically-active cells to pro- or anti-inflammatory phenotypes (*41, 46*). Secretion data from pericytes pre-treated with digoxin and lanatoside C were consistent with an inhibitory effect on IL-1β-induced soluble adhesion molecule (sICAM-1 and sVCAM-1) and CX3CL1 secretion, whilst IL-6 was increased and interkeukin-8 (IL-8) displayed no change (Fig 2 A-L, concentration response curves (fig. S2A)). Because complications often arise with excessive dosage of digoxin, several outputs of pericyte viability following digoxin or lanatoside C treatment were examined to rule out the possibility of the inflammatory-modulating effects being a result of cytotoxicity. While both digoxin and lanatoside C reduced pericyte number, this is likely a reflection of reduced proliferation over the treatment period (fig. S2B), as opposed to cell death, which was largely unchanged. Interestingly, a reduction in PDGFRβ expression was observed in response to digoxin and lanatoside C that was consistent with reduced proliferation at this time point. Signalling through the platelet-derived growth factor receptor β (PDGFRβ) pathway in pericytes promotes their proliferation (*47, 48*) suggesting that the anti-proliferative effect of digoxin/lanatoside C may result from this PDGFRβ depletion.

**Fig. 2:**
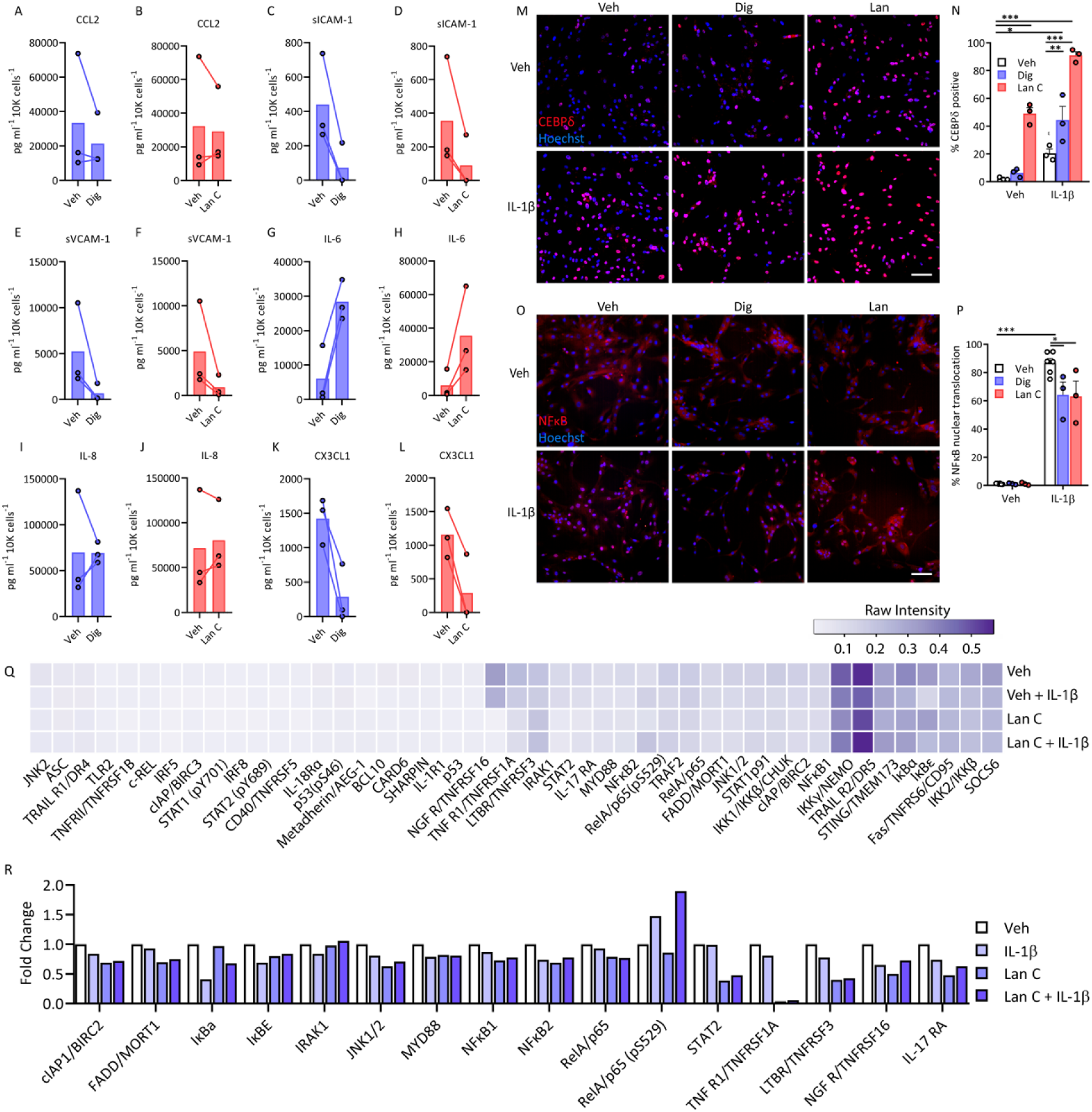
Cardiac glycosides digoxin and lanatoside C modify inflammation-dependent transcriptional responses in pericytes. (A-L) Conditioned media from pericytes after 24 hours of vehicle, digoxin or lanatoside C (125nM points selected from concentration curves in fig. S2) pre-treatment prior to stimulation with IL1β (0.05 ng/mL) for 24 hours (n = 3). (M-P) Pericytes pre-treated with digoxin or lanatoside C for 24 hours were treated with IL-1β (0.05 ng/mL) for 1 hour, and stained for CEBPδ or NFκB, (representative images (M and O, scale = 100 µm), (N) quantification of CEBPδ percent positive cells, and (P) percent NFκB nuclear translocation were compared to control conditions. Two-way ANOVA with Tukey’s multiple comparison test, \*\*\**p*<0.001, \*\**p*<0.01, \**p*<0.05, (n = 3). (Q) Pericytes were pre-treated with vehicle or lanatoside C (1 µM) for 24 hours, then treated with vehicle or IL-1β (0.05 ng/mL) for 30 minutes. Lysates (500 µg per sample) were analysed using the human NFκB array kit. Average intensity values from duplicate spots were normalised to the average intensity of the reference spots on each blot. R) Select targets from NFκB profiler presented as fold change in intensity from vehicle.

Transcription factor activation is a key component of the inflammatory response, in both negative and positive regulatory pathways. CCAAT/enhancer binding protein delta (CEBPδ) has recently been identified as an anti-inflammatory effector in pericytes, downregulating IL-1β-induced CCL2 and ICAM-1 expression (*43*). Conversely, in response to IL-1β, TNFα, and LPS, nuclear factor kappa-light-chain-enhancer of activated B cells (NFκB) translocates to the nucleus where it induces transcription of numerous pro-inflammatory transcripts (*17, 49*). Lanatoside C induced CEBPδ nuclear expression under basal conditions, while both digoxin and lanatoside C increased IL-1β -induced expression (Fig. 2M and N). In contrast both compounds reduced IL-1β-induced NFκB nuclear translocation (Fig. 2O and P) and TGFβ1-induced SMAD2/3 translocation (fig. S3A and B), with no effect on IFNγ-induced signal transducer and activator of transcription 1 (STAT1) activation (fig. S3C and D), suggesting a selective element of inflammatory pathway inhibition. Further inspection of NFκB pathway components from pericyte cell lysates revealed reduced activity of known upstream elements including myeloid differentiation primary response 88 (MyD88), interleukin-1 receptor (IL-1R) associated kinase (IRAK), nuclear factor of kappa light polypeptide gene enhancer in B-cells inhibitor, alpha and epsilon (IκBα, IκBε) and TNF receptor signalling, at the same time specifically increasing phosphorylation of NFκB/p65 at S529 (Fig. 2 Q and R). Interestingly, gene expression analysis of pericytes under the above conditions do not show a diminished inflammatory response-quite the opposite. Both digoxin and lanatoside C increased transcripts for CCL2, ICAM-1, VCAM-1, Il-6, IL-8, CX3CL1 and CEBPδ (fig. S4) suggesting that these compounds may be acting on multiple pathways to reduce inflammatory responses.

In mixed glial cultures, while neither digoxin or lanatoside C reduced IL-1β-induced NFκB translocation in microglia (Fig. 3A to D), cytokine secretions were consistent with pericyte cultures (Fig. 3E to M). This is most likely due to the low percentage of microglia present in the predominantly pericyte cultures at early passage (7.261 %, ± 4.709 PU.1 positive cells). This suggests that digoxin and lanatoside C do not modulate NFκB translocation-dependent inflammatory effects in human microglia.

**Fig. 3:**
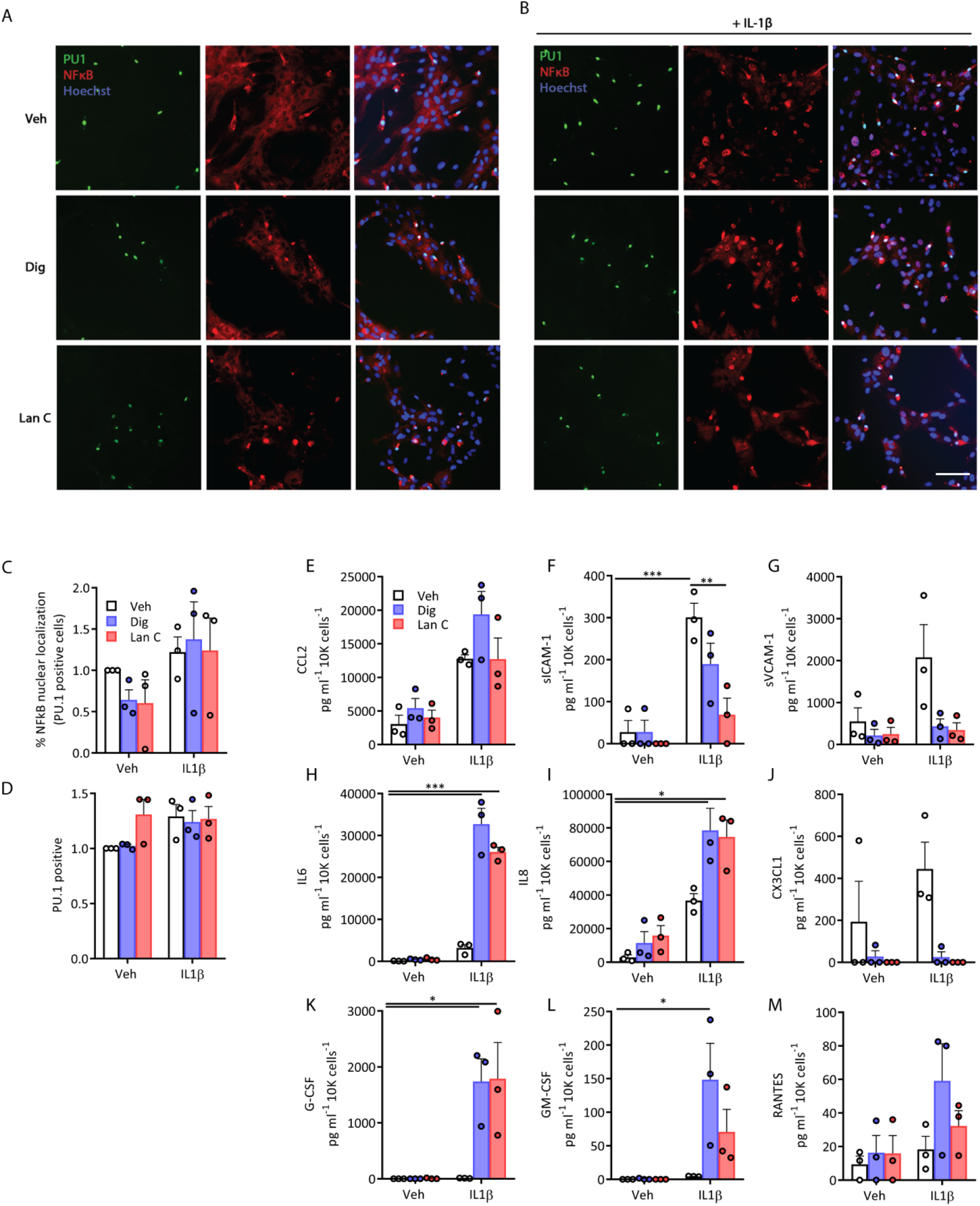
Cardiac glycosides modulate cytokine secretions from mixed glial cultures of microglia and pericytes. (A, B) Mixed glial cultures were treated with cardiac glycosides for 24 hours prior to stimulation with IL1β for 1 hour (A representative images). Inflammatory response was assessed by NFκB nuclear translocation in PU.1 positive microglia (C, D). (E-M) Conditioned media from mixed glial cultures after 24 hours of vehicle, digoxin (100 nM) or lanatoside C (1 µM) pretreatment prior to stimulation with IL1β (0.05 ng/mL) for 24 hours (n = 3). Two-way ANOVA with Tukey’s multiple comparison test, \*\*\**p*<0.001, \*\**p*<0.01, \**p*<0.05.

### Anti-inflammatory effects of digoxin and lanatoside C in primary human brain endothelia

Brain endothelia lining the cerebral blood vessels are the first point of contact for therapeutic drugs in the systemic circulation and express numerous active efflux transporters which complicates CNS drug delivery. Additionally, brain endothelia play an important role in neuroinflammatory responses generated from both the periphery and the brain parenchyma by mediating recruitment, adhesion, and transcellular/paracellular immune cell trafficking across the BBB through expression of cellular adhesion molecules (ICAM-1, VCAM-1) and chemokines (IL-8, CCL2) (*50-52*). As p-glycoprotein substrates, an intact BBB would likely exclude digoxin and lanatoside C entrance to parenchymal brain cells, although this barrier is often weakened during inflammatory states or neurological diseases and could facilitate cerebral penetrance. In order to further assess anti-inflammatory effects of digoxin and lanatoside C, particularly with respect to leucocyte extravasation in the cell type most likely to be exposed *in vivo*, inflammatory responses of primary adult human brain endothelia were assessed using the aforementioned paradigm described for pericytes.

Expression of endothelial markers Claudin-5, ETS-related gene (ERG), zonula occludens-1 (ZO-1), and platelet/endothelial cell adhesion molecule-1 (PECAM/CD31) were confirmed in primary cultures of human brain endothelia *in vitro* (Figure 4A to D). While neither digoxin nor lanatoside C significantly altered basal CCL2 or ICAM-1 expression by ICC, digoxin significantly increased CEBPδ expression (Fig. 4E to I). Both digoxin and lanatoside C blocked IL-1β-induced secretion of all analytes measured by CBA (CCL2, sICAM-1, sVCAM-1, IL-6, IL-8, CX3CL1, chemokine(C-C motif) ligand 5/regulated on activation, normal T cell expressed and secreted (RANTES), granulocyte colony-stimulating factor (G-CSF), and granulocyte-macrophage colony-stimulating factor (GM-CSF) (Figure 3J to R).

**Fig. 4:**
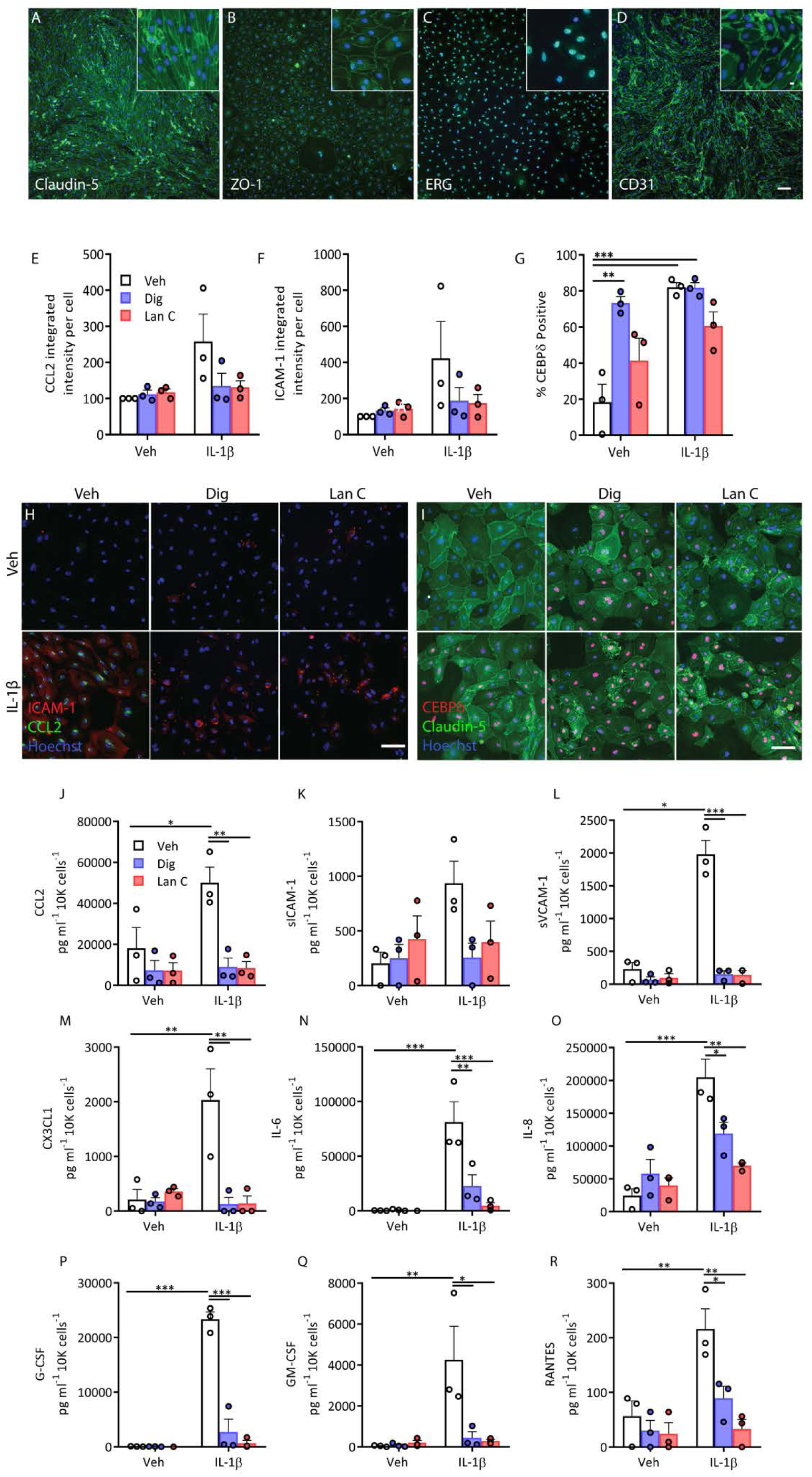
Inflammatory cytokine secretion is blocked by both digoxin and lanatoside C in primary human brain endothelial cells. Endothelial cells derived from human brain tissue express endothelial markers, Claudin-5 (A), ZO-1, (B), ERG (C), and CD31 (D). (E-G) Human endothelial cells were pretreated with digoxin (100 nM) or lanatoside C (1 µM) for 24 hours, followed by 24 hours of IL-1β (0.05 ng/mL) treatment. Staining quantification of CCL2 (E), ICAM-1 (F), and nuclear CEBPδ (G), in endothelial cells. Representative images of (n = 3) (H, I). (J-R) Conditioned media was analysed by CBA in endothelial cells treated as above (n = 3). Cytokine/chemokine concentration was normalized to total cell counts (Hoechst)). Two-way ANOVA with Tukey’s multiple comparison test \*\*\**p*<0.001, \*\**p*<0.01, \**p*<0.05.

### Human meninges and choroid plexus as 3-D *ex vivo* targets

Leptomeninges envelop the brain and spinal cord and are composed of the arachnoid and pia mater. The subarachnoid space within the leptomeninges is bathed in CP-derived CSF and plays a crucial role in protecting the CNS. The anatomical relationship of the vascular cells, meninges, and CP through the subarachnoid, Virchow-robin/perivascular space, and the CSF-containing ventricles allows for the exchange of fluid components be it therapeutics or less desired inflammatory constituents (*53*). However, the meninges and CP can also contribute to numerous neurological disorders, due to their intimate interactions and exposure to the peripheral environment respectively through the meningeal lymphatics and the fenestrated blood vessels (*29, 54-56*). Early immune responses occur in multiple sclerosis, and meningitis in the meninges and CP as both are sites of immune cell trafficking into the CNS (*27, 57-59*). Further, inflammatory secretions in the subarachnoid space in the ventricular space by the CP can penetrate the brain to alter cerebral functioning, likely as a consequence of glymphatic influx, whereby neurovascular cells including pericytes, endothelia, and astrocytes will also be subjected to cytokine presence (*13, 60*). In order to further investigate potential anti-inflammatory functions of digoxin and lanatoside C, we first established a method allowing for the *ex vivo* cultures of human leptomeninges and CP to study inflammation, and potential drug interventions in a culture system more appropriately recapitulating the *in vivo* microenvironment than can be achieved by standard *in vitro* cultures.

Leptomeningeal explants (LME) derived from drug resistant epilepsy biopsy tissue and post-mortem neurologically normal and disease brains (fig. S5 A to D) were found to be viable in culture, even three months after the initial isolation, as determined by live imaging using the ReadyProbes Live/Dead reagents (fig. S5E to G). Immunohistochemical interrogation of cell specific proteins in LME at experimental endpoints revealed mostly vascular cells, staining positively for pericyte markers PDGFRβ, endothelial cells (CD31, *Ulex europaeus* agglutinin I (UEA-1/lectin)), smooth muscle cells (αSMA), fibroblasts (PDZ and LIM domain protein 3 (PDLIM3)) and extracellular matrix components (collagen, type I, alpha I (COL1A1)) (Fig. 5A)(*61*). We also detected cells positive for microglia/macrophage markers Iba-1, HLA-DR and CD68 in both LME and CPE after more than one month in culture (fig S6). Whilst the inflammatory secretome and cytokine-specific responses of brain pericytes and endothelia have been studied to some extent, significantly less is known regarding this response in the leptomeninges. Secretions from LME in response LPS, IL-1β, and IFNγ closely mirrored what we saw in pericytes and endothelia with adhesion molecule and chemokines responsible for immune cell recruitment largely being induced by IL-1β, and LPS, and a lesser response with IFNγ (*22*) (Fig 5B to G, fig S7A). Thus, we used cytokine profiler arrays, which allows for simultaneous detection of 102 different soluble inflammatory mediators in response to the aforementioned immunogenic stimuli (LPS, IL-1β, and IFNγ). As a likely reflection of their cell heterogeneity, the LME demonstrated an extensive basal secretion of inflammatory mediators which was differentially induced by LPS, IL-1β, and IFNγ (Fig. 5H, fig S8A). Importantly, these mediators had no effect on cell viability (fig. S5F). Notably basal detection of several immune mediators was observed from the LME, in particular angiogenin, angiopoetin-2, chitinase 3-like, C-X-C motif chemokine 5 (CXCL5), growth/differentiation factor 15 (GDF-15), insulin-like growth factor protein-2 (IGFBP-2), IL-8, osteopontin, and high basal levels of serpin E1. The most drastic change in secretion in response to cytokine treatment was interferon γ-induced protein-10 (IP-10/CXCL10), which is induced by both IFNγ and LPS, as well as GM-CSF -by IL-1β, and IL-6 by IFNγ and IL-1β.

**Figure 5:**
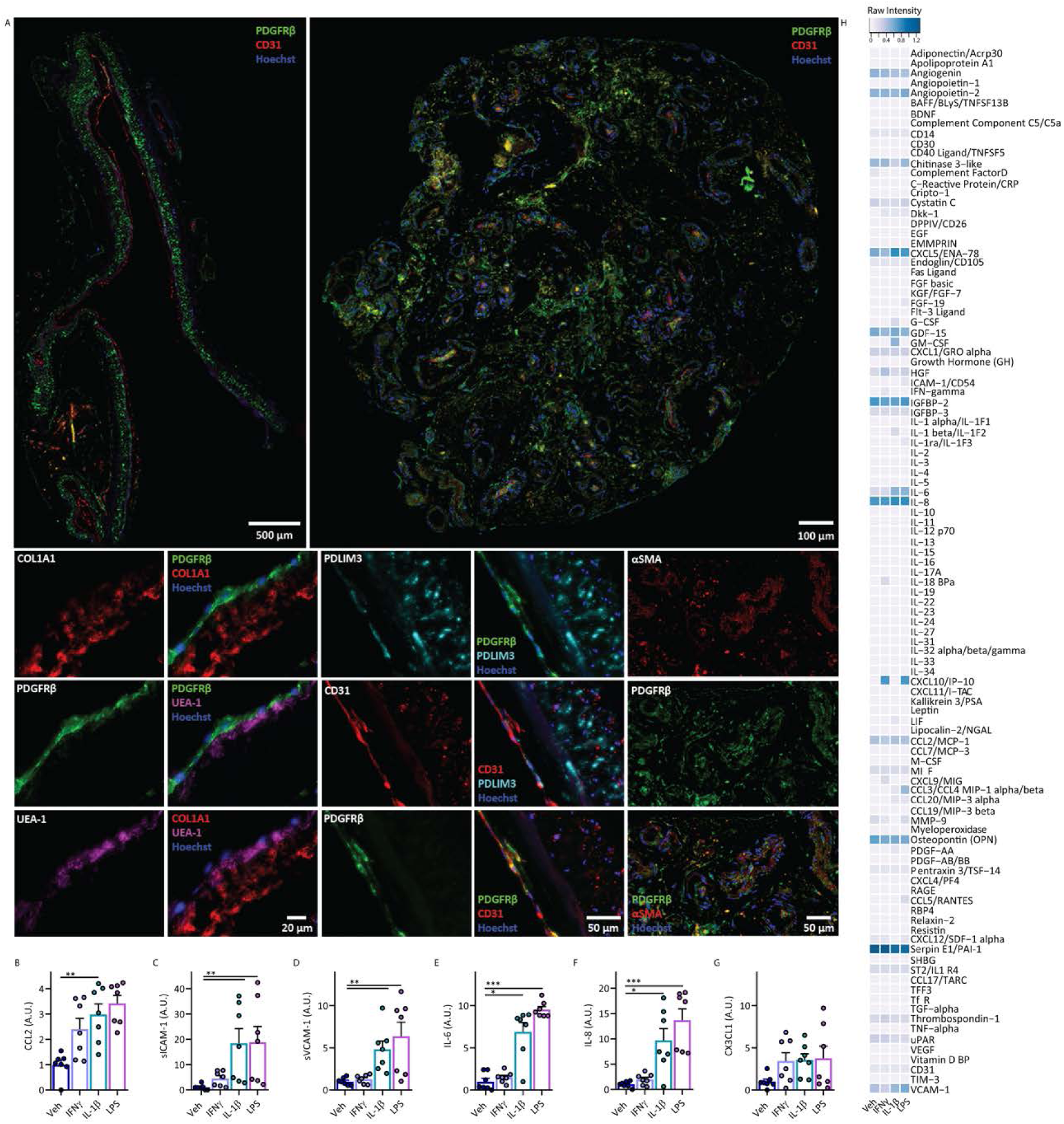
Characterization of leptomeningeal explant cell types and inflammatory signature. A) Immunohistochemical staining of LME for vascular cells types including pericytes (PDGFRβ) endothelial cells (CD31), fibroblasts (PDLIM3), smooth muscle cells (αSMA) and extracellular matrix protein deposition (COL1A1). (B-G) Secretions from meningeal explants following inflammatory stimulation were investigated using cytometric bead arrays (n = 2 cases, 3-4 explants per case), values normalized to vehicle condition, raw values Fig S7, 8. (H) Proteome profilers were used to detect secretions from human brain meningeal explants (pooled from 5 explants per condition, one case) after 24 hours of inflammatory stimulation with vehicle, IFNγ, IL-1β, or LPS (10 ng/mL), raw intensities were normalized to reference spots.

We sought to investigate whether digoxin and lanatoside C could reduce this pro-inflammatory response. As was observed for pericytes and endothelia, digoxin and lanatoside C reduced IL-1β-induced inflammatory mediators in the leptomeninges including CCL2, sICAM-1, sVCAM-1 and CX3CL1 (Fig 6A-I, fig S7B). Using the proteome profilers to interrogate the extent of inhibition by digoxin or lanatoside C we found that both drugs decreased IL-1β –induced secretion of G-CSF, GM-CSF, C-X-C motif chemokine 11 (CXCL11), platelet-derived growth factor AA (PDGF-AA), and sVCAM-1, and they increased secretion of IL-1β dependent IL-6, macrophage inflammatory protein 1-alpha (MIP1α) and TNFα consistently across three LME cases (Fig 6J, fig S7B).

**Figure 6.**
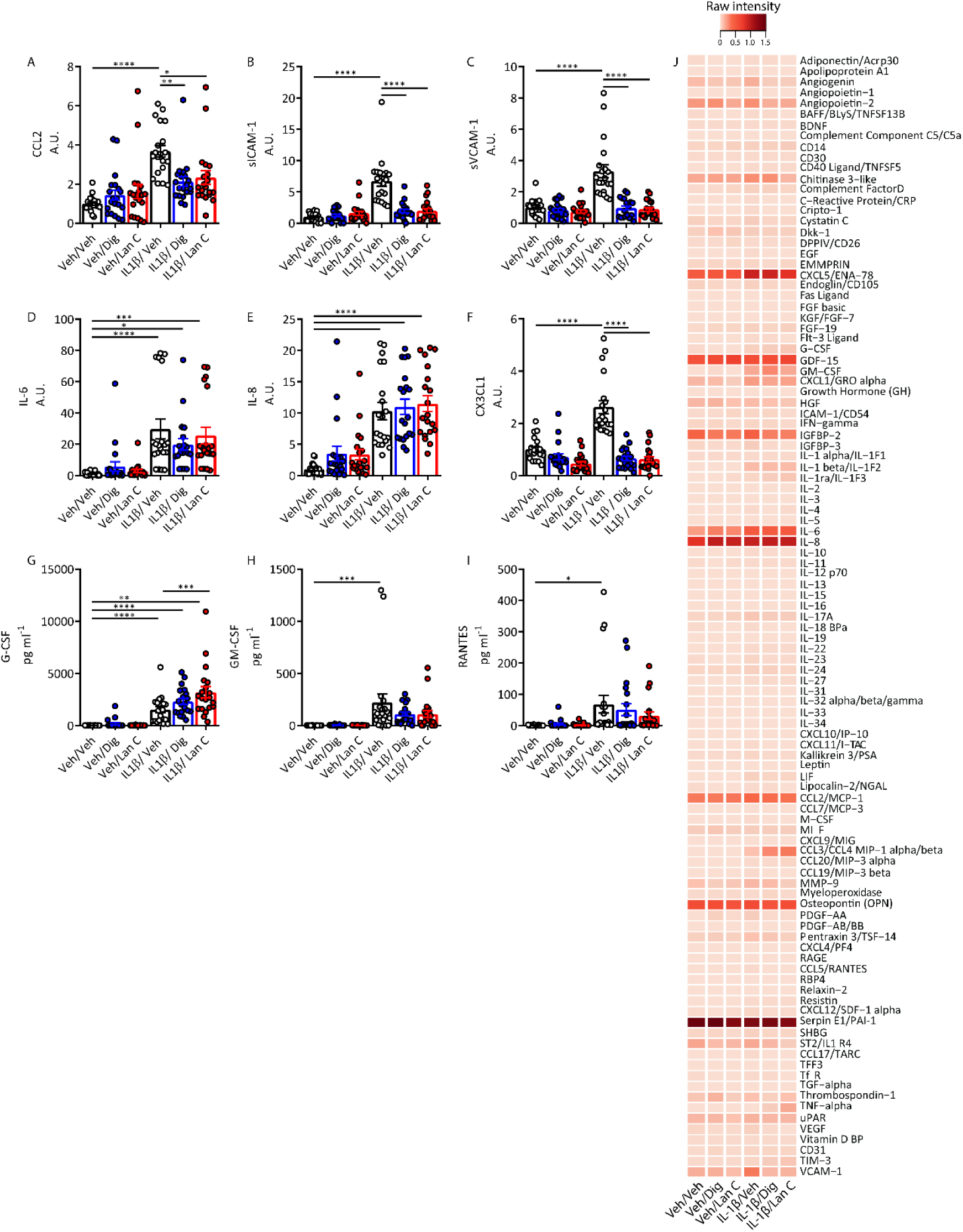
Cardiac glycosides modulate inflammatory responses in leptomeningeal explant cultures. A-I) meningeal explant secretions were measured using CBA (n = 3-5 explants per condition, 5 cases). Secretions in pg/mL were normalized to vehicle for each case-arbitrary units (A.U.) except for G-CSF, GM-CSF and RANTES where cytokines were below the detection level in vehicle conditions. Two-way ANOVA with Tukey’s multiple comparison test \*\*\*\**p*<0.0001, \*\*\**p*< 0.001, \*\**p*< 0.01, \**p*<0.05. J) Proteome profilers were used to characterise LME secretions in response to vehicle or IL-1β (10 ng/mL) with either digoxin or lanatoside C pre-treatment (10 µM). Average intensity values from proteome profilers were normalised to the reference spots on each blot (heatmap is average of n=3 cases, each pooled from 3-4 explants per case).

Using the same paradigm in choroid plexus explants (CPE) we first examined the cellular composition and found the highly vascularized tissue to be harbouring similar cell types to the LME, with the addition of transthyretin-positive epithelia (Fig 7A and B). Similar to the LME, the CPE secreted a range of cytokines in response to pro-inflammatory stimulation albeit to a lesser extent than the LME (Fig 7C-L, fig S8 and S9). Just like the LME under basal conditions, the CPE had undetectable levels of G-CSF, and GM-CSF, very low expression of RANTES, sICAM-1, and noticeable expression of chitinase 3-like, GDF-15, IGFBP-2, IL-8, osteopontin and serpin E1. Expectedly CPE cultures responded to cytokine treatment by changing their secretome. Digoxin and lanatoside C pre-treatment resulted in reduced IL-1β-responses such as inhibition of sICAM-1, sVCAM-1 and CX3CL1 secretion by CBA (Fig 8 A-I). Proteome profilers of CPE conditioned media in contrast to LME show an increase in IL-1β induced secretion of G-CSF, GM-CSF, and CXCL5, but reduced VCAM-1, angiogenin, osteopontin, and interleukin 1 receptor-like 1 (ST2) by digoxin and lanatoside (Fig 8).

**Figure 7:**
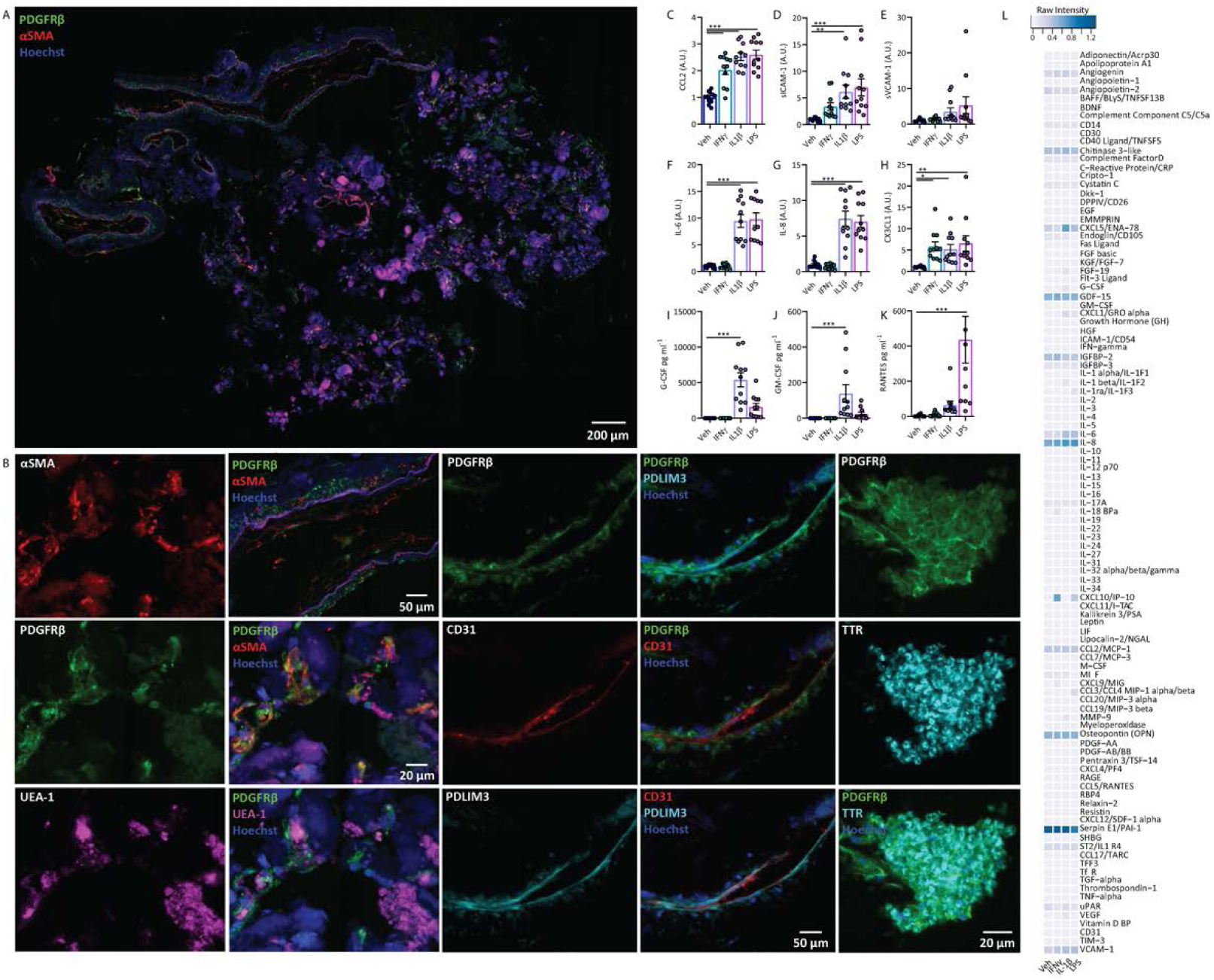
Characterization of choroid plexus explant cell types and inflammatory signature. (A, B) Immunohistochemical staining of CPE for vascular cells types including pericytes (PDGFRβ) endothelial cells (CD31), fibroblasts (PDLIM3), smooth muscle cells (αSMA) and epithelial cells (transthyretin (TTR)). (C-K) Secretions from CPE following inflammatory stimulation were investigated using cytometric bead arrays (n = 3 cases, 3 explants per case), values normalized to vehicle condition, raw values Fig S8, 9. One way ANOVA, Dunnett’s multiple comparisons test, \*\*\**p*< 0.001, \*\**p*< 0.01, \**p*<0.05. (L) Proteome profilers were used to detect secretions from human brain choroid plexus explants (average of n = 2 cases each pooled from 3 explants per condition) after 24 hours of inflammatory stimulation with vehicle, IFNγ, IL-1β, or LPS (10 ng/mL), raw intensities were normalized to reference spots.

**Figure 8.**
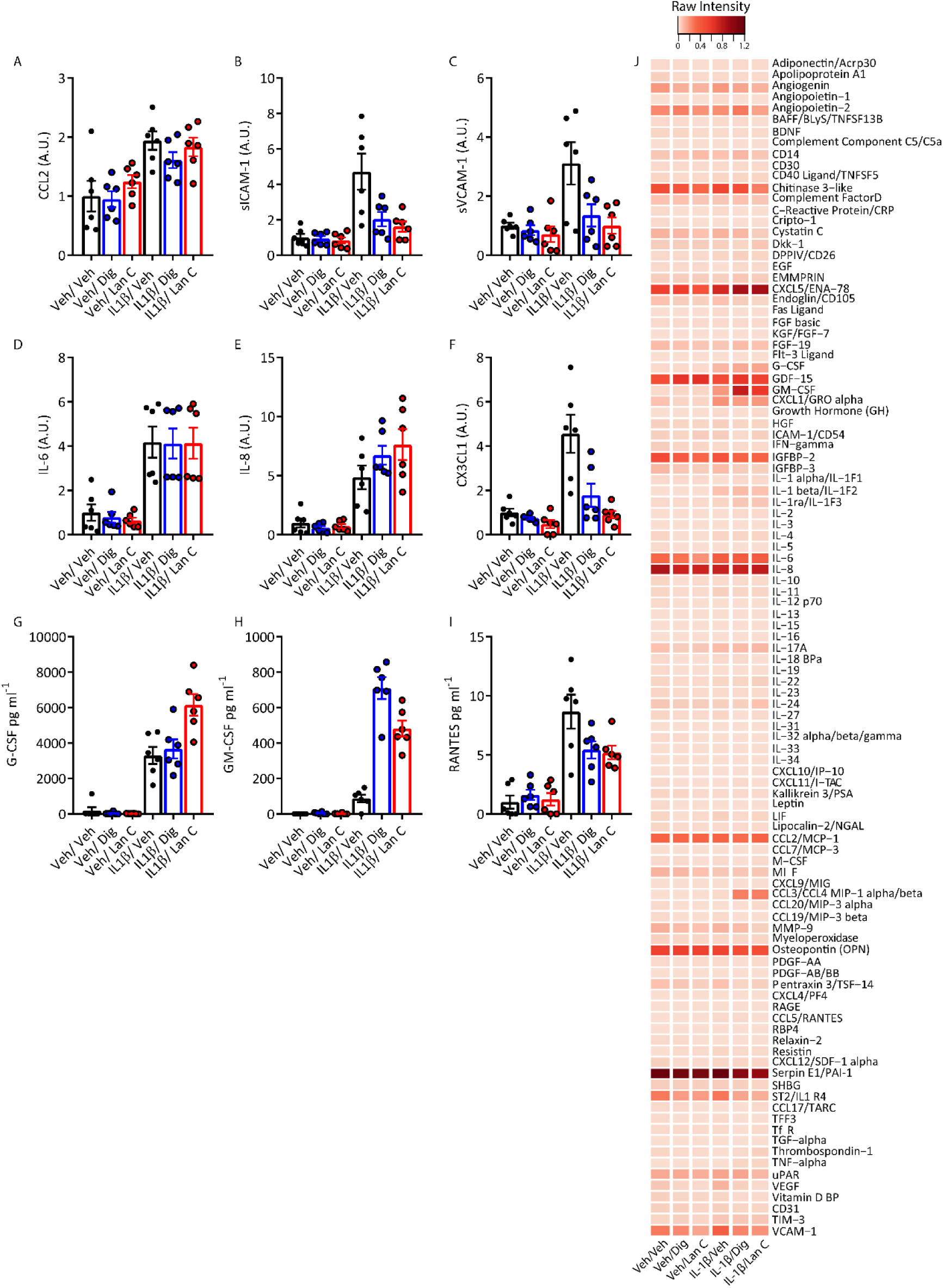
Cardiac glycosides modulate inflammatory responses in choroid plexus explant cultures. A-I) Choroid plexus explants were treated with vehicle, digoxin or lanatoside C (both 10 µM) for 24 hours followed by IL1β (10 ng/mL) for 24 hours. Secretions were measured using CBA (n=2 cases, 3 explants per case). Values normalized to vehicle for each case, except for G-CSF, GM-CSF and RANTES as vehicle condition was under the level of detection, (raw data provided in supplementary, Fig S9). J) Proteome profiler analysis of pooled choroid plexus explant secretions (3 pooled explants per case). Data are presented from the average of each case normalised to reference point.

## Discussion

Neuroinflammation is present in almost every neurological disorder and contributes to disease pathogenesis by precipitating neuronal loss and BBB dysfunction (*62, 63*). As such, the identification of therapeutic compounds that effectively attenuate neuroinflammation has the potential to be beneficial in a diverse range of neurodegenerative diseases, neuropsychiatric disorders, and acute brain injuries. Unfortunately, compounds that were beneficial in preclinical models of neurological disease, including anti-inflammatory therapies, have largely failed to effectively translate to clinical use (*64, 65*). Whilst the failure of anti-inflammatory interventions may be a function of inappropriate timing in the treatment regime, the lack of translational therapeutics could be equally attributed to the frequent use of rodent models in neurological drug discovery, in which responses may poorly reflect those observed in humans. Failure may also be associated with targeting of general inflammatory pathways irrelevant to neurodegenerative disease, suggesting that targeting distinct CNS neuroinflammation pathways may prove more successful.

To address this issue, we utilised primary human brain pericytes to screen for anti-inflammatory drugs in a library consisting of >1280 FDA approved compounds. These cells have the obvious benefit of being of human origin, therefore negating any species differences which complicate observed responses. Additionally, unlike human microglia or astrocytes, these cells are highly proliferative *in vitro*, allowing for efficient bulking and bio-banking of these cultures to generate sufficient yields to undertake large screens (*66*). Due to its ubiquitous presence in neurological disease, we sought to target neuroinflammation in this screen in particular because together, endothelia and pericytes are the major mediators of leucocyte extravasation during neurodegenerative diseases, particularly in AD and MS, through CCL2-directed chemotaxis and ICAM-1 and VCAM-1-mediated adhesion (*52, 67-69*). However, this approach would be equally suited to identify mitogenic enhancers, autophagy regulators, phagocytic inducers or pro-survival drugs, utilising differing stimulation paradigms.

The strength of the current screening approach resides in the fact that compounds underwent several rounds of analysis in primary human brain cells derived from diverse healthy and diseased post mortem and biopsy samples. The drug effects were consistent across different donors, emphasizing the capacity of this screening platform as a tool to identify modifiers of human-specific targets taking into account the inherent human variation from the first stages of drug discovery. With recent advancements in the ability to isolate and culture cells from the human brain, in addition to deriving these cells from patient induced pluripotent stem cells, these screening approaches can be expanded to human microglia, astrocytes, and even neurons (*70-72*).

Whilst the use of non-human or immortalized cell lines has potential issues with such high-throughput screens, so too does the use of *in vitro* primary cultures themselves. Pericytes and endothelia utilised here were subjected to several rounds of passaging in order to obtain sufficient yields, for which phenotypic drift can occur (*73, 74*). It is therefore unclear whether cells accurately reflect the phenotype observed *in vivo*. In order to analyse the anti-inflammatory effects of digoxin and lanatoside C in a more complex, multi-cellular model without the complications of dissociated and passaged cells, a protocol to simply and effectively isolate leptomeningeal and choroid plexus explant cultures was used (*75*). These explants display excellent viability, they are easily accessible from the post-mortem brain, or from neurosurgical specimens and their ease of handling makes them an attractive model. The advancements in organotypic slice cultures could soon allow for coculture methods of LME and CPE with human neurons to define cellular interactions in disease conditions (*76*). The explants displayed an extensive inflammatory secretome, including cytokines, chemokines, and adhesion molecules. Of equal importance, the anti-inflammatory effects observed using *in vitro* monocultures of pericytes and endothelial cells were largely consistent with our *ex vivo* leptomeningeal and CPE cultures, suggesting the appropriateness of both model systems in identifying immune modulating compounds. The ability of these compounds to attenuate inflammatory responses in several distinct cell types is promising with the ability to effectively attenuate global neuroinflammation, as this response is not restricted to one particular region. Furthermore, neither digoxin nor lanatoside C completely blocked inflammatory responses, as can be deleterious, with a basal level of inflammatory capabilities important for homeostasis and defence responses (*77*).

The utilisation of an FDA-approved compound library is advantageous as compounds have already undergone safety trials in humans, easing the repurposing of drugs for other disorders. As such, potential information about efficacy of such compounds can often be gleaned from retrospective analysis of patient cohorts taking these compounds for other indications, therefore expensive and timely clinical human safety trials are not necessary. Further, extensive data are already available on their pharmacological properties (*45, 72, 78*). Digoxin and lanatoside C are structurally very similar and well-known for their inhibitory effects on the Na^+^/K^+^-ATPase pump (*79*). Both act similarly in cancer cell lines to inhibit proliferation having actions on tumour necrosis factor-related apoptosis-inducing ligand (TRAIL), Src and protein kinase C δ (PKCδ) pathways. Digoxin is approximately three times more potent than lanatoside C, which is consistent with the data presented here (*45, 80-82*). Currently, evidence for digoxin and lanatoside C in anti-inflammatory mechanisms is limited, with suggested effects on leukocyte infiltration and extravasation through inhibition of cytokine/chemokine secretion and reduced NFκB expression (*83*). Although the efficacy of other cardiac glycosides as anti-inflammatory compounds in brain barrier tissues has not been investigated, cardiac glycosides demonstrate a large range of chemical diversity and absorption, distribution, metabolism, elimination and toxicity (ADMET) properties (*79*). Nonetheless, we have shown that these compounds can act on multiple inflammatory pathways in brain barrier cells including pericytes, endothelia, microglia, and in meningeal and choroid plexus explants. Therefore they have strong potential as anti-inflammatory drugs in the context of neuroinflammation.

Neurodegenerative diseases are often a result of several diverging dysregulated pathways, including inflammatory responses, protein aggregation, BBB breakdown, and inappropriate clearance mechanisms. As such, the identification of multi-target, multi-action drugs (targeting both neuroinflammatory pathways and blood-brain barrier cells) will prove more effective than single target therapeutics (*7*). Moreover, accessibility to vascular cells via perivascular spaces, and tissues such as the meninges and CP through the brain boundary regions and CSF compartments should be considered in drug delivery strategies. Here we demonstrate the utility of repurposed drug screens in human brain cells and identify drugs for use as therapeutic agents in attacking the neurodegenerative cascade in cells and tissues that can be more easily reached. Taken together, our described pipeline represents a promising approach for neuroinflammatory and neurodegenerative drug development.

## Materials and Methods

### Study Design

This study was designed to identify chemical compounds that could modulate inflammatory responses in human brain vascular cells. Using post-mortem brain tissue of healthy and disease patients, as well as epilepsy biopsy tissue we screened 1280 FDA-approved compounds against an IL-1β-induced inflammatory response. Candidates identified from the initial screen by CCL2 and ICAM-1 expression in pericytes were forwarded for secretion analysis, and transcription factor activity in pericytes, endothelial cells, and mixed glial cultures. Meningeal and choroid plexus explants were tested as an *ex vivo* model of neuroinflammation.

### Tissue source

Biopsy human brain tissue was obtained from the Neurological Foundation Human Brain Bank, with informed written consent, from the middle temporal gyrus (MTG) of patients undergoing surgery for drug-resistant epilepsy. Post-mortem leptomeninges were obtained from regions overlying the MTG, and CPE were derived from the lateral ventricle from one hemisphere from neurologically normal individuals, or those with various neurological diseases (Table S1). All brain tissue collection and processing protocols were approved by the Northern Regional Ethics Committee, (AKL/88/025/AM09 New Zealand) for biopsy tissue, and the University of Auckland Human Participants Ethics Committee (Ref # 011654, New Zealand) for the post-mortem brain tissue. All methods were carried out in accordance with the approved guidelines.

### Primary human brain mixed glial cultures

Biopsy human brain tissue was obtained from the MTG of patients with drug-resistant epilepsy and mixed glial cultures, containing microglia, astrocytes, and pericytes, were generated as described previously (*71*). Mixed glial cultures were maintained in complete media (DMEM/F12 with 10% FBS and 1% PSG (penicillin 100 U/mL, streptomycin 100 µg/mL, L-glutamine 0.29 mg/mL)) at 37 °C with 5% CO_2_ until confluent. Flasks were trypsinized with 0.25% Trypsin-1mM EDTA and scraped to detach firmly adherent microglia. Viable cells were counted based on trypan blue exclusion and 5,000 cells/well were seeded into 96-well plates in complete media and used for experimentation after 1-3 days. All experiments performed on mixed glial cultures were at passage two.

### Primary human brain pericyte culture

To generate pure pericyte cultures, mixed glial cultures were sub-cultured up to passage four in order to eliminate non-proliferative microglia and astrocytes as described previously (*71, 84*). Pericyte cultures were maintained in complete media at 37 °C with 5% CO_2_. Viable cells were counted based on trypan blue exclusion and 5,000 cells/well were seeded into 96-well plates in complete media and used for experimentation after 1-3 days. All experiments performed on pericytes were at passages 4-9.

### Primary human brain endothelial culture

Primary human endothelial cells were isolated from brain microvessels as described previously (*22*). Endothelial cultures were maintained in Endothelia Cell Media (ECM; ScienCell) at 37 °C with 5% CO_2._ Viable cells were counted based on trypan blue exclusion and 10,000 cells/well were seeded into 96-well plates in complete media and used for experimentation when cells had produced a 100% confluent monolayer (typically around five days). All experiments performed on endothelia were at passages three-five.

### Compound screening

High-throughput screening to identify compounds which modulate inflammatory responses in pericytes was performed using the Prestwick Chemical Library (Prestwick Chemical, France) containing 1280 small molecules of FDA approved drugs. Controls included vehicle for compounds (DMSO), TGFβ_1_ (1 or 10 ng/mL), vehicle for IL-1β (0.01% BSA in PBS), or media alone. Pericytes were pre-treated with compounds (or controls) in duplicate at 10 µM for 24 hours before treatment with IL-1β (0.05 ng/mL) for another 24 hours. At endpoint cells were fixed and immunostained for CCL2 and ICAM-1 and nuclei were counterstained with Hoechst 33258 (as detailed below). Cells were imaged using the ImageXpress^®^ Micro XLS automated fluorescent microscope and total Hoechst positive cell counts and the integrated intensity per cell of CCL2 and ICAM-1 was quantified as described previously (*17*).

### Immunocytochemistry and high-throughput image analysis

At endpoint cells were fixed in 4% paraformaldehyde for 15 minutes and washed/permeabilized with phosphate buffered saline (PBS) with 0.2% Triton X-100™ (PBS-T). Cells were incubated with primary antibodies diluted in goat immunobuffer (1% goat or donkey serum, 0.2% Triton X-100™, and 0.04% thiomersal in PBS) overnight at 4 °C, (dilutions of antibodies are listed in Table S2). Cells were washed again in PBS-T and incubated with fluorescently conjugated secondary antibodies for two hours at room temperature. Nuclei were counterstained by a 15 minute incubation with 20 μM Hoechst 33258 (Sigma). Quantitative analysis of intensity measures and scoring of positively stained cells was performed using the Cell Scoring, Multiwavelength Cell Scoring, and Nuclear Translocation modules on MetaXpress^®^ software (Molecular Devices) as previously described (*48*).

### Cytometric bead array

Conditioned media was collected from samples and centrifuged at 160 ×g for five minutes to collect possible cells and debris. The supernatant was obtained and stored at 20 °C. Analyte concentrations were measured by cytometric bead array (CBA; BD Biosciences, CA, USA) as described previously (*85*). CBA samples were run on an Accuri C6 flow-cytometer (BD Biosciences, CA, USA). Data were analysed using FCAP-array software (version 3.1) (BD Biosciences, CA, USA) to convert fluorescent intensity values to concentrations utilising a 11-point standard curve (0-10,000 pg/mL). Concentrations were normalised to total cells as determined by Hoechst positive counts for monolayer cells.

### Human NFκB pathway array kit

Human pericytes were seeded into 10 cm dishes and allowed to reach confluency. Pericytes were treated with DMSO (0.01%) or lanatoside C (0.1 µM) for 24 hours then treated with IL-1β (0.05 ng/mL) for 30 minutes. Cells were lysed on ice with kit lysis buffer containing protease inhibitor tablets (Roche), scraped into pre-chilled tubes and incubated on ice for 30 minutes with intermittent vortexing. Lysates were centrifuged at 14,000 × g for 15 minutes, and protein concentration of the supernatant was quantified using the DC protein quantification kit (Bio-Rad). Samples were stored at -80 °C until array analysis. Arrays were carried out following the manufacturer’s instructions, with 500 µg of protein from each sample used for analysis. Membranes were imaged using the LI-COR Odyssey^®^ FC imaging system (LI-COR Biosciences). Intensity of individual spots was determined using Image Studio™ Lite and values were normalized to the average of the six reference spots on each blot.

### Cytokine profiler

Conditioned media from 3-6 leptomeningeal explants was pooled, centrifuged at 160 × g for five minutes, and the supernatant was collected and stored at -20 °C. Secretome analysis of the clarified media was performed using the Proteome Profiler™ Human XL Cytokine Array Kit (R & D Systems), as per manufacturer’s instructions. Chemiluminescent detection of the membranes was performed on the LI-COR Odyssey^®^ FC (LI-COR Biosciences). Intensity of individual spots was determined using Image Studio™ Lite and values were normalized to the average of the six reference spots on each blot.

### EdU proliferation assay

Proliferation of pericytes was determined by incorporation of the thymine analogue 5-ethynyl-2’-deoxyuridine (EdU) with the Click-iT^®^Assay Kit (Life Technologies C10340) according to manufacturer’s instructions. Briefly, EdU (5 µM) was added to cells 24 hours prior to endpoint and incubated at 37 °C. Cells were fixed with 4% PFA for 15 minutes at room temperature, rinsed with 3% BSA in PBS, and permeabilized with 0.5% Triton X-100 in PBS for 20 minutes at room temperature. Cells were washed twice with 3% BSA in PBS and then EdU reaction cocktail was added for 30 minutes at room temperature protected from light. Cells were then washed once more with 3% BSA in PBS and imaged as described above.

### Primary human leptomeningeal and choroid plexus explant culture

Leptomeninges were removed by gross dissection from the brain overlying the MTG, from autopsy tissue of neurologically normal or pathologically confirmed cases of neurodegenerative disorders (AD, PD, HD, and FTD), as previously described (*86*). Leptomeningeal tissue was washed in complete media in sterile petri dishes and dissected into pieces ∼ 2 mm^3^. Leptomeningeal explants were placed in individual wells in 500 μL of complete media in a 24-well plate allowing them to remain in suspension. Media changes were performed twice a week. Explants that failed to alter media acidification, as determined by a phenol red colour change, were deemed non-viable and discarded. Explants were cultured for at least a week before using for experiments to allow equilibration but remained viable for up to four months after initial isolation. For functional studies, individual explants were placed in 100 μL of media within a 96-well plate.

### Explant viability and histological processing

Cell death was determined using the ReadyProbes™ cell viability imaging kit (Invitrogen). NucBlue live reagent and NucGreen dead reagent stains were diluted into media (2 drops per mL blue, 1 drop per mL green), and added to cells in culture 30 minutes prior to endpoint and incubated at 37 °C for 30 minutes. Z-stacks (200-250 slices, 5 μm step) of explants were then acquired using the ImageXpress high content imaging system (Molecular Devices). 2-D projections were used for analysis of positively labelled nuclei with the Custom Module Editor (image analysis pipeline described in detail in supplementary data (fig. S5).

Immunohistochemical staining on formalin fixed, paraffin embedded explants, sectioned at 7µm was done as previously described (*87*). Antigen retrieval with TRIS-EDTA (pH 9) was performed, followed by blocking with 10% donkey serum prior to antibody incubation. Sections were incubated with primary antibodies (for dilutions see Table S2) overnight at 4 °C, rinsed in PBS then incubated with secondary antibodies and Hoechst 33258 (SIGMA) for 3 hours at room temperature in the dark. Sections were imaged using the 20X objective on the Metasystems V-slide scanning microscope and stitched to generate images of whole explants. Control sections where the primary antibody was omitted showed no immunoreactivity. The control experiments showed that the secondary antibodies did not cross-react with each other. All confocal recordings were done using an FV1000 confocal microscope (Olympus) with a 40X oil immersion lens (NA 1.00).

### Statistical Analysis

All cell culture experiments were performed three separate times with cells from different donors. Data were normalized to vehicle controls were indicated in figure legends. Statistical test were performed using Graphpad Prism software, one-way, or two-way ANOVA with Tukey’s *post hoc* analysis. Data are presented as the mean ± SEM.

## Supplementary Materials

Fig. S1. Controls from compound screen and follow-up screen.

Fig. S2. NFκB pathway profiler original blots.

Fig. S3. Concentration response curves for cytokine secretion and viability from pericytes treated with digoxin or lanatoside C (raw data from three cases).

Fig. S4. Gene expression data from pericytes treated with IL-1β and digoxin or lanatoside C.

Fig. S5. Using explant cultures derived from human meninges and choroid plexus for drug testing.

Fig. S6. Microglia/macrophage markers in leptomeningeal and choroid plexus explants.

Fig. S7. Raw data from CBA analysis of leptomeningeal explant secretions.

Fig. S8. Human cytokine proteome profiler original blots and select hits comparison.

Fig. S9. Raw data from CBA analysis choroid plexus secretions

## Acknowledgments

We would like to thank the donors for their generous gift of brain tissue for research. We also thank staff at Auckland Hospital and Marika Eszes (Technical Officer at the Neurological Foundation Human Brain Bank). This work was supported by a Programme Grant from the Health Research Council of New Zealand, the Sir Thomas and Lady Duncan Trust, the Coker Charitable Trust and the Hugh Green Foundation.

## Figures

**Table S1.** Human tissue sources used in this study.

| Case | Tissue source | Confirmed diagnosis | Age | Sex | PMD (hr) | Experiment |
| --- | --- | --- | --- | --- | --- | --- |
| E203 | MTG | Temporal lobe epilepsy | 46 | F | <1 | ML |
| E204 | MTG | Temporal lobe epilepsy | 45 | F | <1 | ML |
| E206 | MTG | Temporal lobe epilepsy | 29 | F | <1 | ML |
| E208 | MTG | Temporal lobe epilepsy | 52 | F | <1 | ML |
| E211 | MTG | Temporal lobe epilepsy | 51 | F | <1 | ML |
| E213 | MTG | Temporal lobe epilepsy | 23 | M | <1 | ML |
| E214 | MTG | Temporal lobe epilepsy | 35 | F | <1 | ML |
| E215 | MTG | Temporal lobe epilepsy | 29 | F | <1 | ML |
| E216 | MTG | Temporal lobe epilepsy | 38 | M | <1 | ML |
| SS37 | TL | Paediatric epilepsy | 9 | M | <1 | ML |
| SS42 | TL | Paediatric epilepsy | 8 | M | <1 | ML |
| SS48 | TL | Paediatric epilepsy | 10 | F | <1 | ML |
| SS50 | TL | Paediatric epilepsy | 3 | F | <1 | ML |
| SS54 | TL | Paediatric epilepsy | 25w | M | <1 | ML |
| H238 | MTG, meninges | normal | 63 | F | 16 | ML, LME |
| H239 | MTG, meninges | normal | 64 | M | 15.5 | ML, LME |
| H250 | meninges | normal | 93 | F | 19 | LME |
| H253 | meninges | normal | 96 | M | 24 | LME, CPE |
| FTD6 | Meninges, CP | Frontotemporal dementia | 76 | M | 7 | LME, CPE |
| HC166 | Meninges, CP | Huntington's disease | 84 | F | 22.5 | LME, CPE |
| HC171 | Meninges, CP | Huntington's disease | 51 | M | 24 | LME, CPE |
| PD85 | meninges/CP | Parkinson's disease | 73 | M | 4 | LME, CPE |
| AZ129 | meninges | Alzheimer's disease | 81 | M | 4.5 | LME |
| AZ133 | meninges/CP | Alzheimer's disease | 96 | M | 21 | LME, CPE |

MTG-middle temporal gyrus, ML-monolayer cultures, CP-choroid plexus, LME-leptomeningeal explants, CPE-choroid plexus explants

**Table S2.** Antibodies and stains used in this study.

| Antibody (species) | Catalogue Number (Source) | Dilution |
| --- | --- | --- |
| αSMA (mouse) | IS611 (Dako) | 1/100 IHC |
| CCL2 (MCP-1) (Rabbit) | ab-9669 (Abcam) | 1/500 ICC |
| CD31 (mouse) | M0823 (Dako) | 1/500 ICC, 1/100 IHC |
| CD68 (KP1)(mouse) | Ab-955(Abcam) | 1/500 IHC |
| CEBPδ (mouse) | sc-365546 (Santa Cruz) | 1/500 ICC |
| Claudin-5 (rabbit) | ab131259 (Abcam) | 1/5000 ICC |
| COL1A1 (mouse) | sc-293182 (Santa Cruz) | 1/50 IHC |
| HLA-DR (mouse) | M0775 (Dako) | 1/1000 IHC |
| IBA1 (goat) | ab-5076 (Abcam) | 1/500 IHC |
| ICAM-1 (mouse) | sc-107 (Santa Cruz) | 1/500 ICC |
| IL-6 (goat) | AF-206 (R&D Systems) | 1/2000 ICC |
| KCNJ8/Kir6.1 (rabbit) | PA5-56628 | 1/100 IHC |
| NFκBp65 (rabbit) | sc-107 (Santa Cruz) | 1/500 ICC |
| PDGFRβ (goat) | AF-385 (R&D Systems) | 1/2000 ICC |
|  |  | 1/200 IHC |
| PDGFRβ (rabbit) | 3169 (Cell Signalling) | 1/100 IHC |
| PDLIM3 (rabbit) | PA5-52099 (Thermo Fisher Scientific) | 1/50 IHC |
| PU.1 (rabbit) | 2258 (Cell Signalling) | 1/500 ICC |
| SMAD2/3 (mouse) | sc-133098 (Santa Cruz) | 1/500 ICC |
| STAT1 (rabbit) | 14994 (Cell Signalling) | 1/500 ICC |
| Streptavidin Alexa Fluor 647 | S21374 (Invitrogen) | 2 mg/mL (1/500) |
| Transthyretin /Pre-albumin (rabbit) | A0002 (Dako) | 1/1000 |
| ZO-1 (mouse) | 339100 (Invitrogen) | 1/500 ICC |
| Lectin (UEA-1) | L8262 (SIGMA) | 1/1000 IHC |
| Hoechst 33258 | SIGMA | 1/500 ICC<br>1/10000 IHC |

957 IHC-immunohistochemistry, ICC-immunocytochemistry

