## Supplementary Information for "Recycling old drugs: cardiac glycosides as therapeutics to target barrier inflammation of the vasculature, meninges and choroid plexus"

Supplementary Material

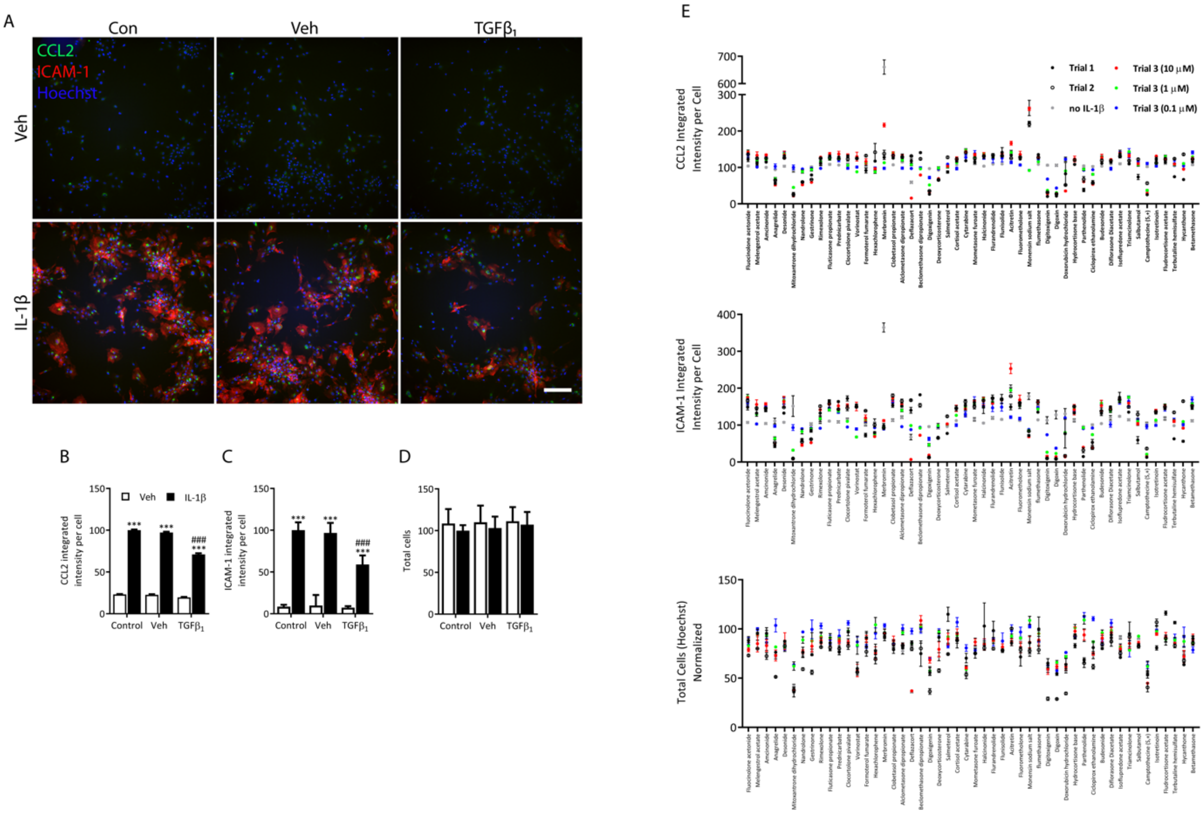


Fig. S1. Consistency of hits and controls across drug screen trials in pericytes. (A-D) Validation of controls used for drug screen shows TGFβ$_1$ (1 ng/mL), a known inhibitor of CCL2 and ICAM-1 expression in pericytes is consistent across all drug plates. Control is untreated, vehicle is DMSO for compounds, or 0.1% BSA in PBS for TGFβ$_1$ . Two-way ANOVA with Tukey's multiple comparison test,****p*<0.001 from control/veh, $^{###}$ *p*<0.001 from control/IL-1β (n = 32 plates).  E) Potential drug hits from three trials (including trial 3, at three concentrations, and one trial with no IL-1β treatment). Immunocytochemical analysis of CCL2, ICAM-1 and total cell counts, normalized to vehicle treatment.

A

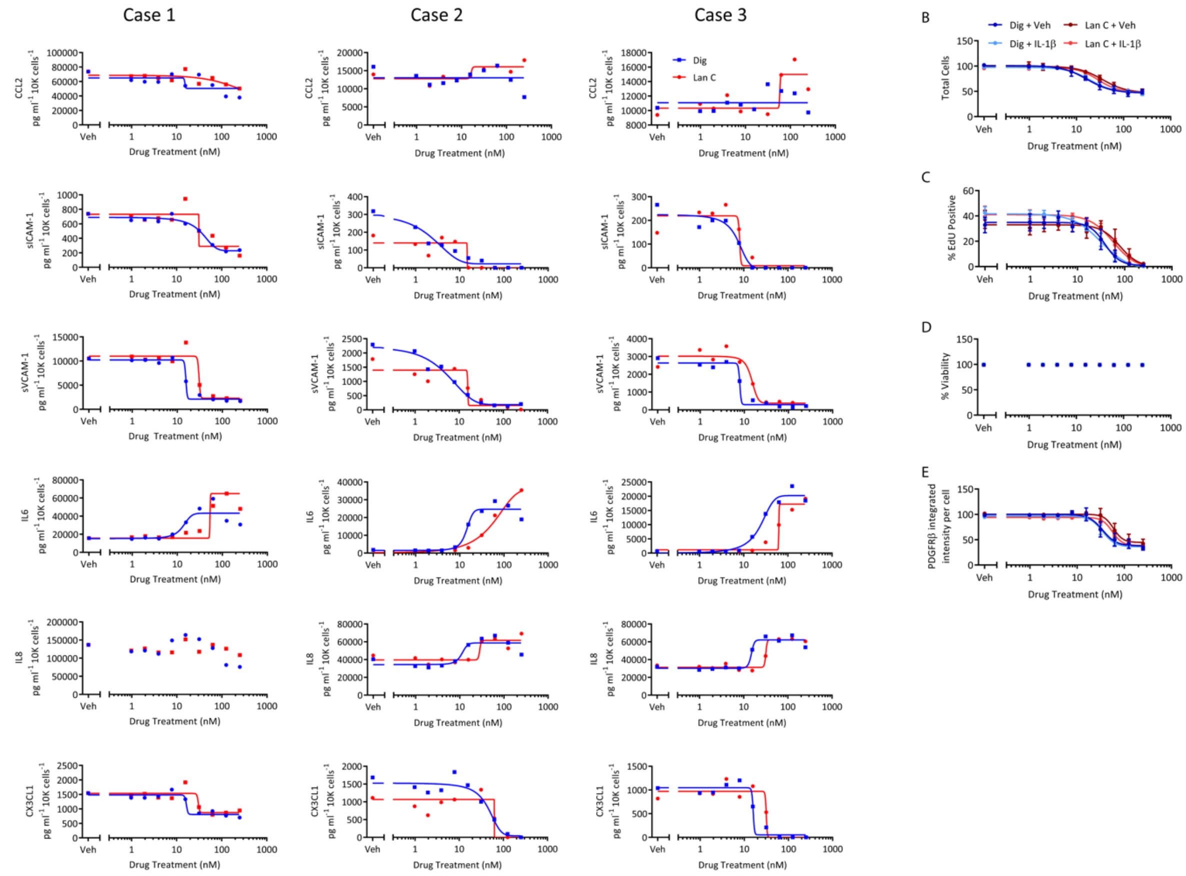

Fig. S2.

Concentration response of pericytes to digoxin or lanatoside C in the presence of IL-1β.

A) Pericytes were pretreated with digoxin or lanatoside for 24 hours prior to IL-1β stimulation. Secretions were quantified with cytometric bead array and normalized to total cells. B) Cells treated as above were assessed for proliferation (EdU), viability (LDH), and PDGFRβ expression.

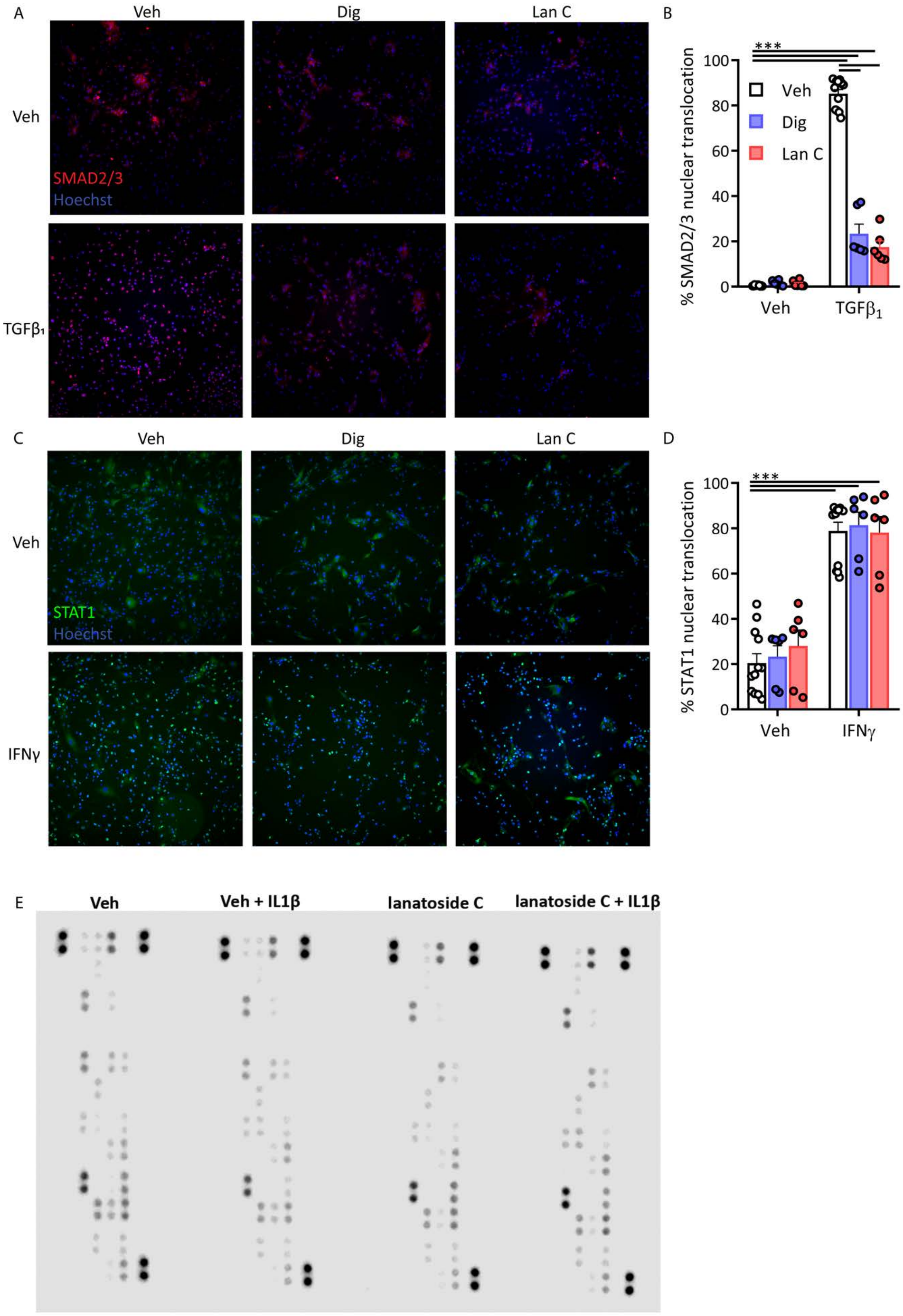

A
Veh
Dig
Lan C
Veh
SMAD2/3
Hoechst
TGFβ₁
B
% SMAD2/3 nuclear translocation
***
Veh
Dig
Lan C
Veh
TGFβ1
C
Veh
Dig
Lan C
Veh
STAT1
Hoechst
IFNγ
D
% STAT1 nuclear translocation
***
Veh
IFNγ
E
Veh
Veh + IL1β
lanatoside C
lanatoside C + IL1β

Fig. S3. Pericytes pre-treated with digoxin (0.1 $\mu$M) or lanatoside C (1 $\mu$M) for 24 hours were treated with TGF$\beta_1$ (1 ng/mL) for one hour and stained for SMAD2/3 (A) or IFN$\gamma$ (1 ng/mL) for one hour and stained for STAT1 (B). Quantification of nuclear translocation of SMAD 2/3 (C) or STAT1 (D), ***$p$<0.001, (n=3).(E) Blots from NF$\kappa$B profiler arrays, quantified in figure 2.

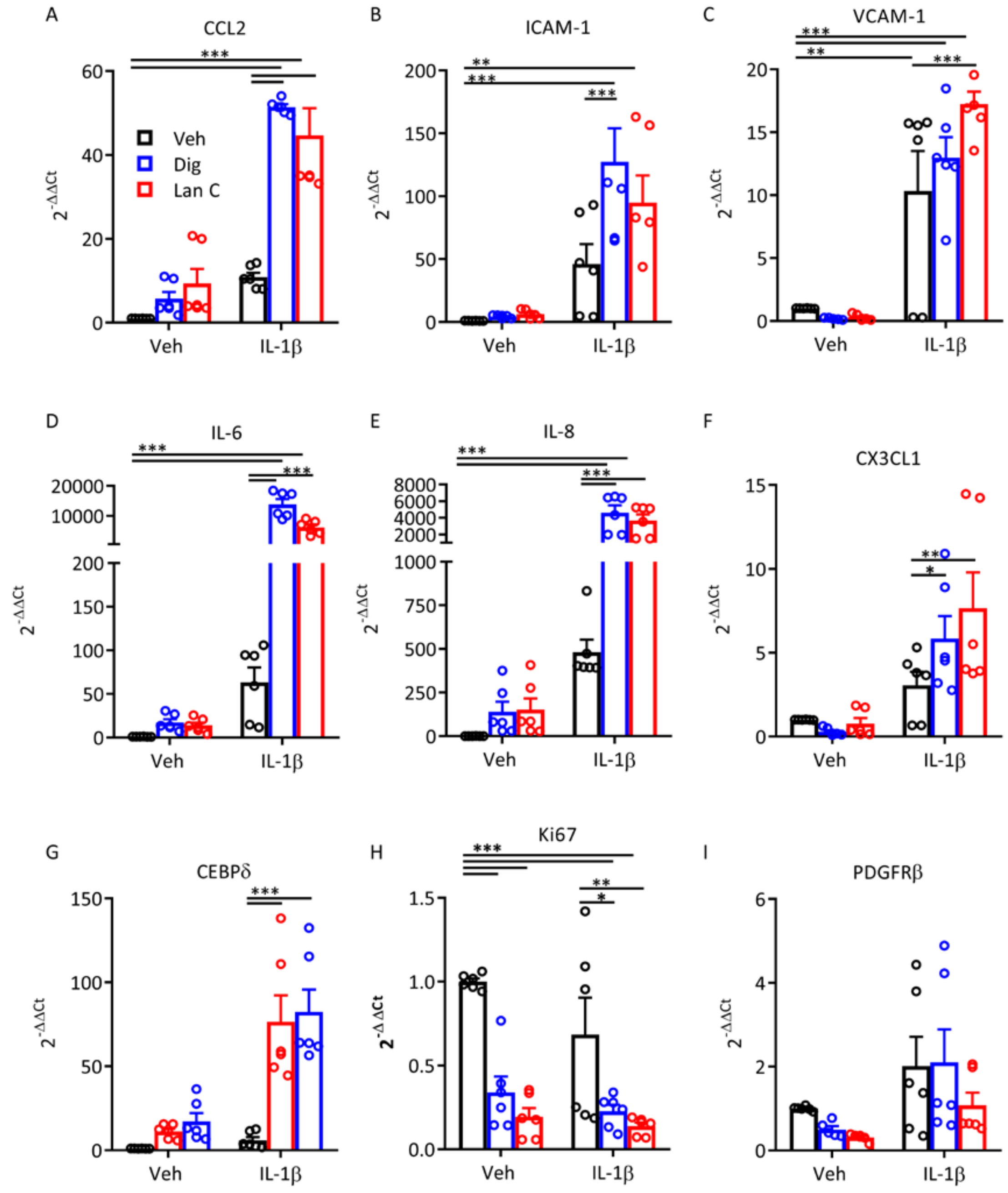


Fig S4. Cardiac glycosides do not block transcriptional activation of IL-1β-dependent inflammatory responses in pericytes. Cells were treated for 24 hours with digoxin (0.1 μM) or lanatoside C (1 μM) then treated with IL-1β for 24 hours. Statistical analysis performed using two-way ANOVA, ***$p$<0.001, **$p$<0.01, *$p$<0.05 (n = 3 cases, 2 experimental replicates).

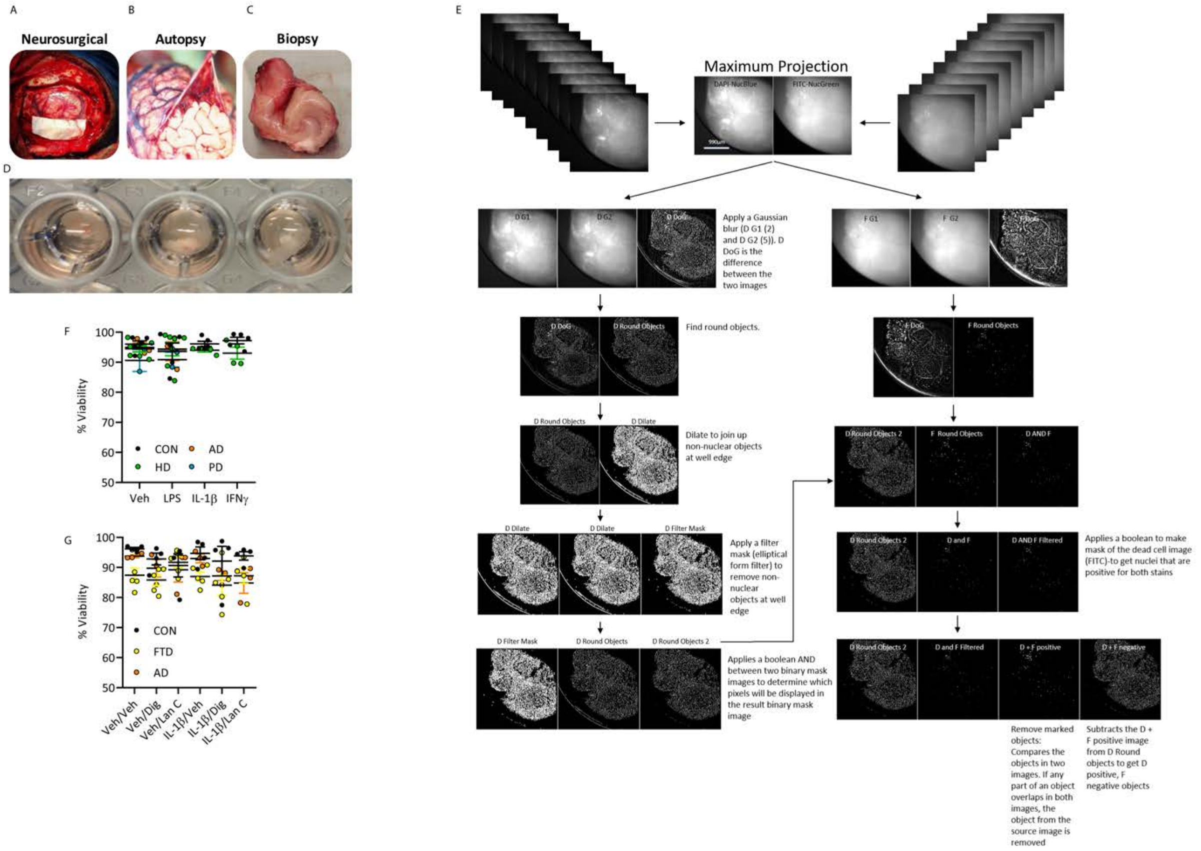


Fig S5. Development of leptomeningeal/choroid plexus explant cultures for ex vivo pharmacological studies. A-C) meninges are sourced from neurosurgeries for the treatment of epilepsy, autopsy or tumour biopsies. Tissue is diced into 2 mm2 and placed into cell culture dishes with DMEM/F12, 10% FBS, 1% PSG (penicillin 100U/ml, streptomycin 100 µg/ml, L-glutamine 0.29 mg/ml)) at 37 °C with 5% $CO_2$. E) Viability analysis was performed using the custom module editor function of MetaXpress software to quantify live and dead cells from Z-stacks of explants. F-G) Quantification of explants after treatment with cytokines for 24 hours (10 ng/mL), or with pretreatment of digoxin or lanatoside C (10 µM) for 24 hours, followed by IL-1β (10 ng/mL) for 24 hours.

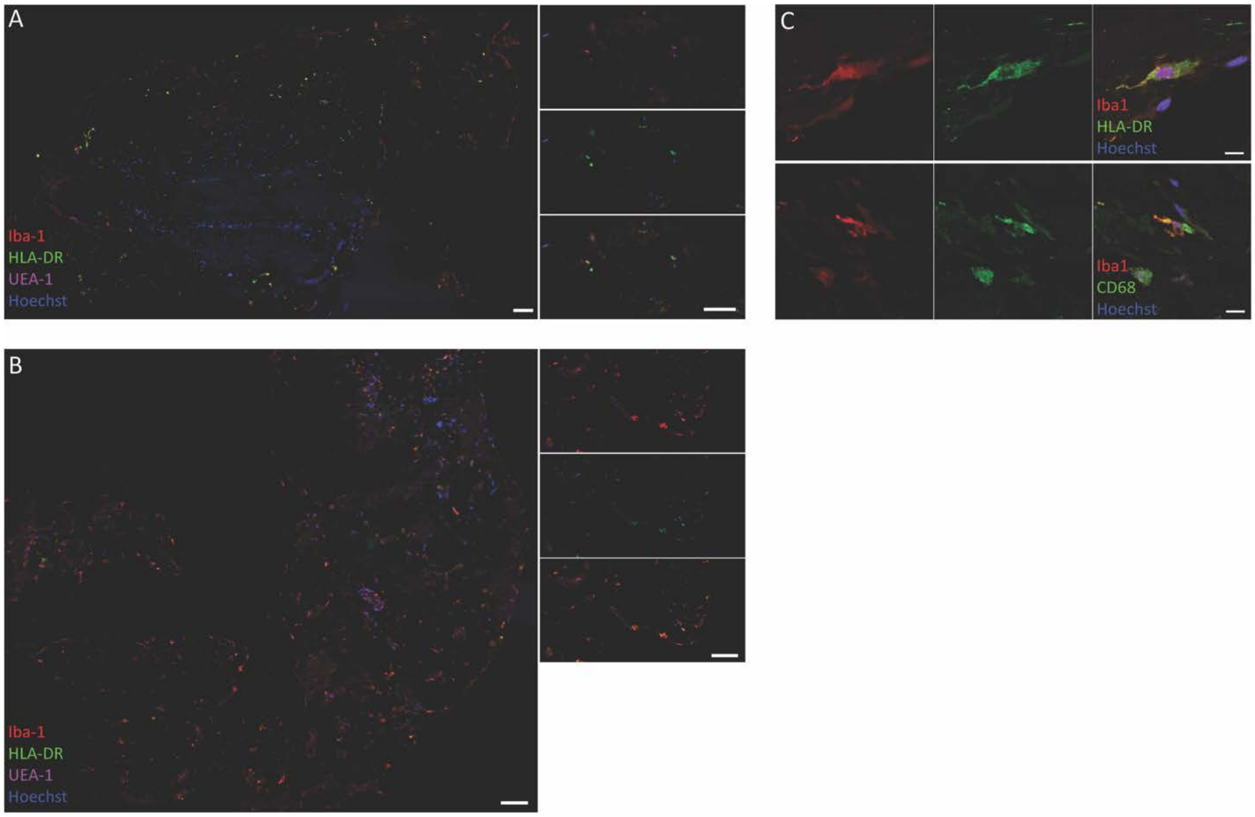


Fig. S6. Microglia/macrophage presence in leptomeningeal and choroid plexus explants. Immunohistochemical staining of microglial/ macrophage markers in leptomeningeal explants (A) (scale 100 μm, and in choroid plexus explants (B) (scale = 100 μm (left) and 50 μm (right). C) confocal images demonstrating co-expression of Iba-1 with HLA-DR and CD68 (scale = 10 μm).

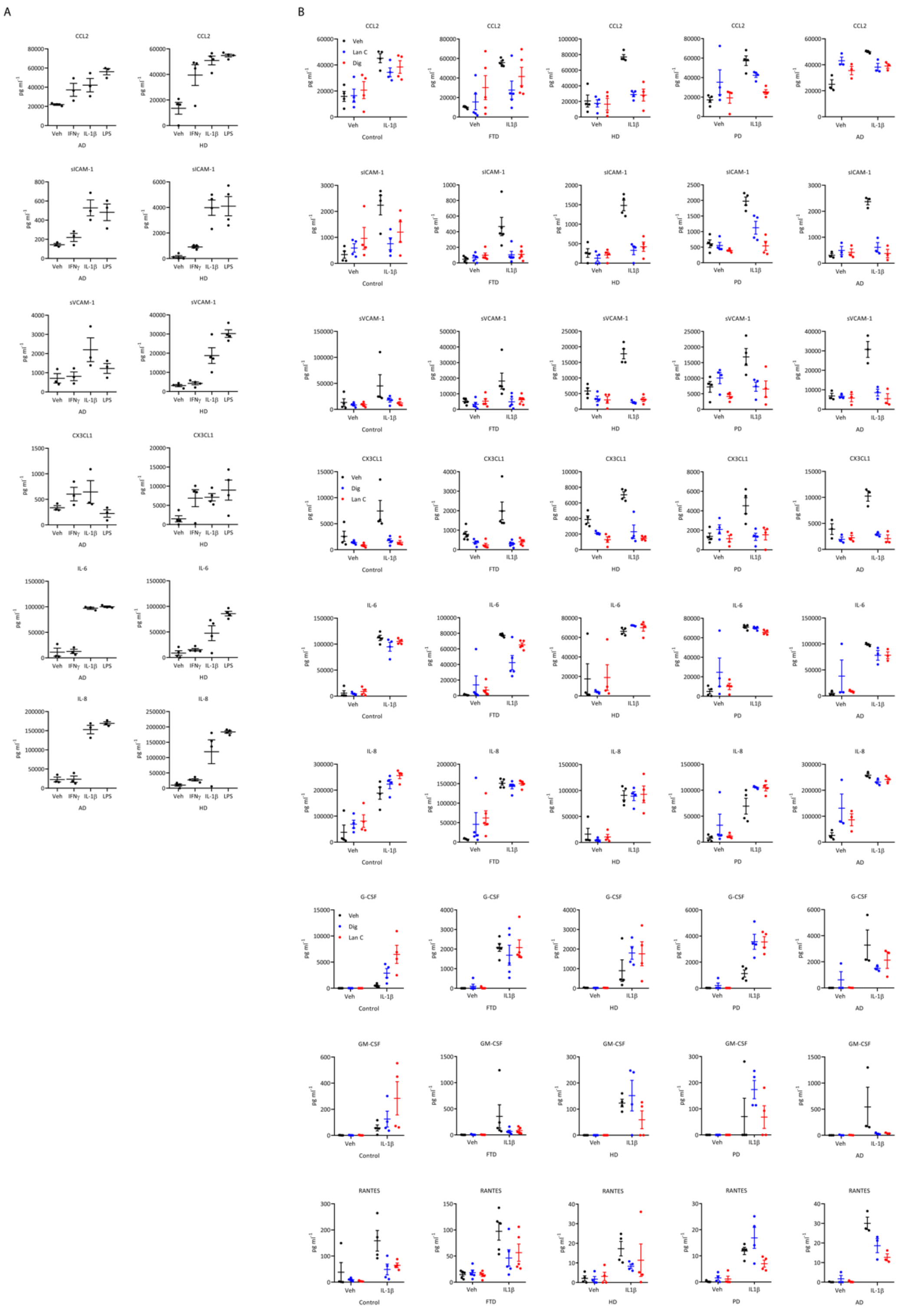


Fig S7. A) Raw cytokine concentration from meningeal explants from two different donors treated with vehicle, IFNγ, IL-1β, or LPS (10 ng/mL), for 24 hours. B) Raw cytokine concentrations from meningeal explants from five different donors LME cases treated with digoxin or lanatoside C (10 μM) for 24 hours then IL-1β (10 ng/mL) for 24 hours. Each point is a single explant. (Control- neurologically normal, FTD- frontotemporal dementia, HD-Huntington's disease, PD-Parkinson's disease, AD-Alzheimer's disease).

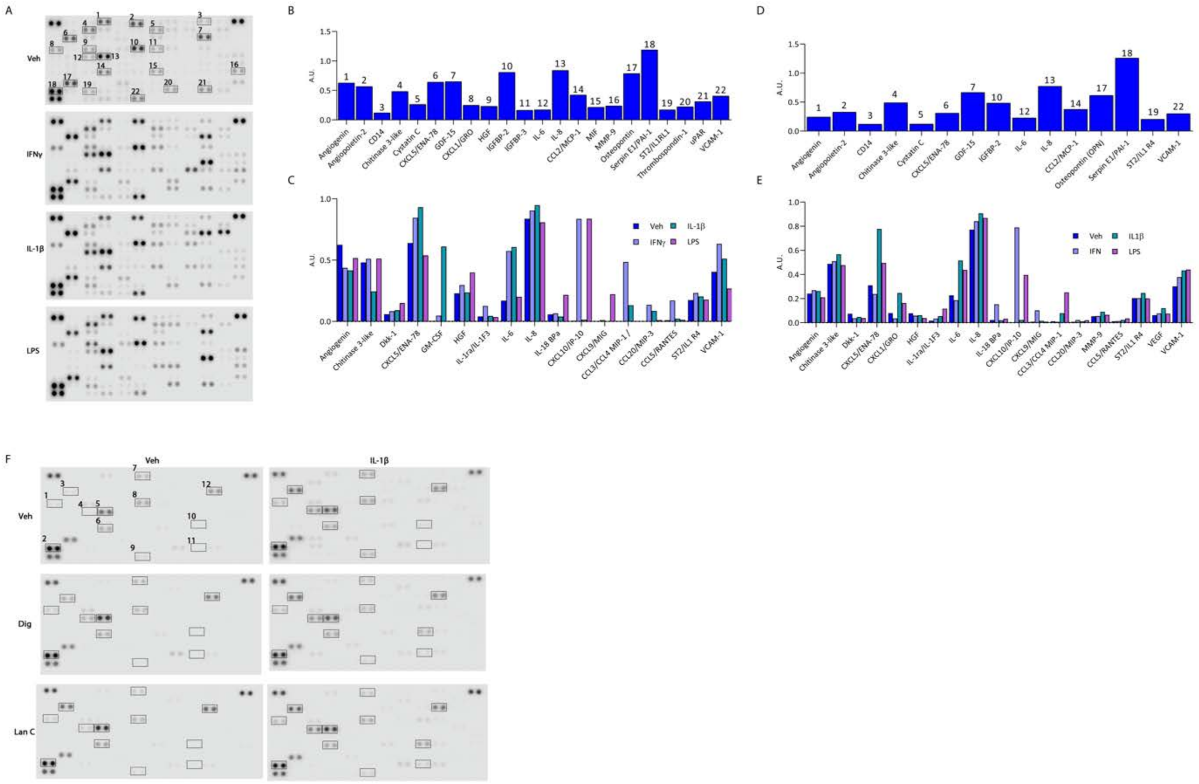


Fig S8. Human cytokine proteome profiler original blots and select hits comparison.

A) Original blots of LME conditioned media exposed to vehicle (Veh), IFNγ, IL-1β, or LPS after 24 hours. Normalized intensity values above an arbitrary threshold under basal conditions in LME (B) or CPE (D). Select secreted proteins that are modified by cytokine treatment in LME, or D, E) in CPE. F) Proteome profilers were used to characterise the IL-1β (10 ng/mL) induced changes in secretions from meningeal explants in response with either digoxin or lanatoside pre-treatment (10 µM), representative blots from one case, pooled secretions from 5 explants. (1-CXCL1, 2- SerpinE1, 3-CXCL5, 4-IL-6, 5-IL-8, 6-CCL2, 7-Ang-2, 8-IGFBP2, 9-VCAM-1, 10-MIP1α, 11-TNFα, 12-GDF-15).

A

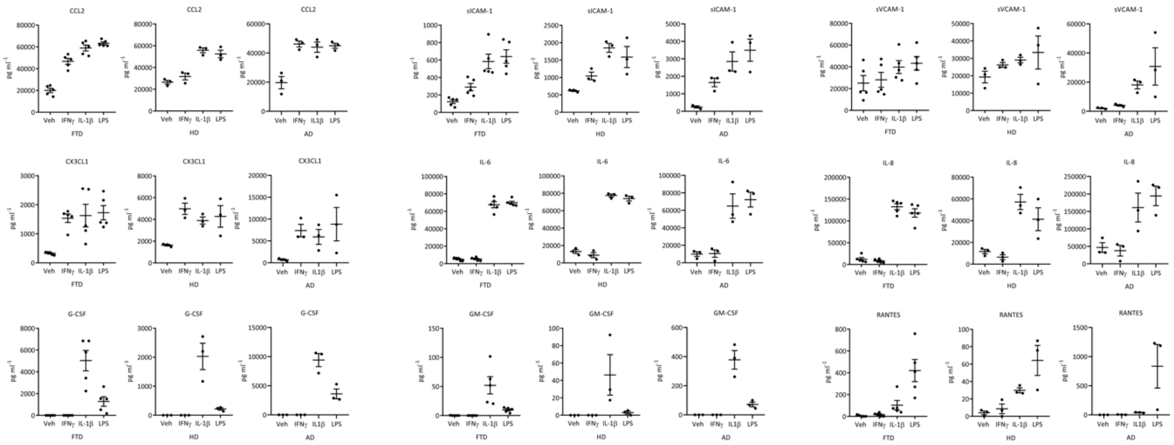


B

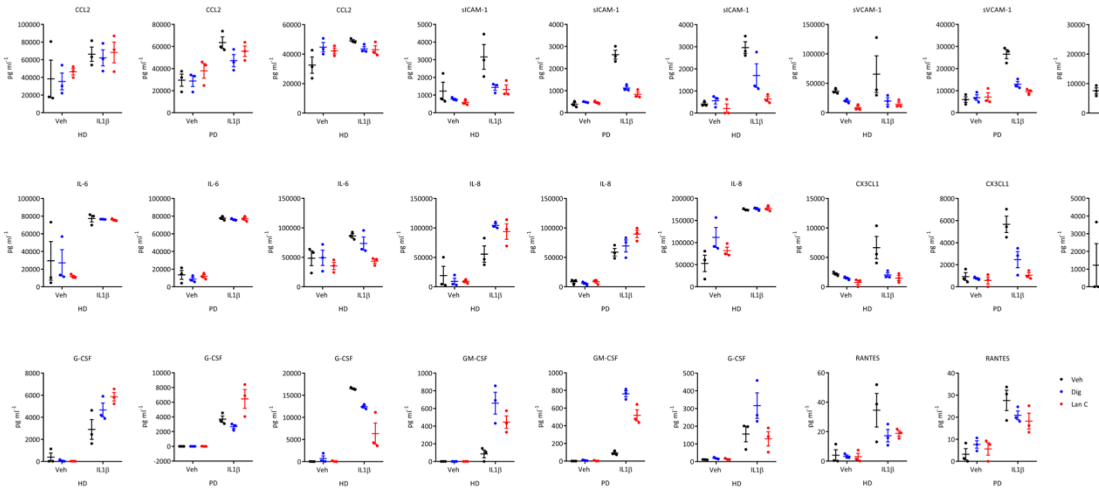


Fig. S9: Raw CBA data from analysis of choroid plexus secretions.
A) Raw cytokine concentrations from choroid plexus explants from three individual donors (n=3 explants per case) that were treated with vehicle, IFNγ, IL-1β or LPS (10 ng/mL), for 24 hours. B) Choroid plexus explants from three individual donors (3 explants per case) were treated with vehicle, digoxin or lanatoside C (10 µM) for 24 hrs, then vehicle or IL-1β (10 ng/mL) for 24 hours. (FTD-frontotemporal dementia, HD-Huntington's disease, PD-Parkinson's disease, AD-Alzheimer's disease).
